# “Nitrogen demand, supply, and acquisition strategy control plant responses to elevated CO_2_ at different scales”

**DOI:** 10.1101/2023.11.30.567584

**Authors:** Evan A. Perkowski, Ezinwanne Ezekannagha, Nicholas G. Smith

## Abstract

Plants respond to elevated atmospheric CO_2_ concentrations by reducing leaf nitrogen content and photosynthetic capacity – patterns that correspond with increased net photosynthesis rates, total leaf area, and total biomass. Nitrogen supply has been hypothesized to be the primary factor controlling these responses, as nitrogen availability limits net primary productivity globally. Recent work using evo-evolutionary optimality theory suggests that leaf photosynthetic responses to elevated CO_2_ are independent of nitrogen supply and are instead driven by leaf nitrogen demand to build and maintain photosynthetic enzymes, which optimizes resource allocation to photosynthetic capacity and maximizes allocation to growth. Here, *Glycine max* L. (Merr) seedlings were grown under two CO_2_ concentrations, with and without inoculation with *Bradyrhizobium japonicum*, and across nine soil nitrogen fertilization treatments in a full-factorial growth chamber experiment to reconcile the role of nitrogen supply and demand on leaf and whole-plant responses to elevated CO_2_. After seven weeks, elevated CO_2_ increased net photosynthesis rates despite reduced leaf nitrogen content and maximum rates of Ribulose-1,5-bisphosphate (RuBP) carboxylase/oxygenase (Rubisco) carboxylation and electron transport for RuBP regeneration. Effects of elevated CO_2_ on net photosynthesis and indices of photosynthetic capacity were independent of nitrogen fertilization and inoculation. However, increasing nitrogen fertilization enhanced positive effects of elevated CO_2_ on total leaf area and total biomass due to increased nitrogen uptake and reduced carbon costs to acquire nitrogen. Whole-plant responses to elevated CO_2_ were not modified by inoculation across the nitrogen fertilization gradient, as plant investment toward symbiotic nitrogen fixation was similar between CO_2_ treatments. These results indicate that leaf nitrogen demand to build and maintain photosynthetic enzymes drives leaf photosynthetic responses to elevated CO_2_, while nitrogen supply regulates whole-plant responses. Our findings build on previous work suggesting that terrestrial biosphere models may improve simulations of photosynthetic processes under future novel environments by adopting optimality principles.

## Introduction

Terrestrial ecosystems are regulated by complex carbon and nitrogen cycles. As a result, terrestrial biosphere models, which are beginning to include coupled carbon and nitrogen cycles (Shi *et al*., 2016; Davies-Barnard *et al*., 2020; Braghiere *et al*., 2022), must accurately represent these cycles under different environmental scenarios to reliably simulate carbon and nitrogen fluxes (Oreskes *et al*., 1994; Prentice *et al*., 2015). While the inclusion of coupled carbon and nitrogen cycles in terrestrial biosphere models was intended to improve model reliability, large uncertainty in the role of nitrogen availability and nitrogen acquisition strategy on leaf and whole plant responses to increasing atmospheric CO_2_ concentrations persists (Arora *et al*., 2020; Davies-Barnard *et al*., 2020; Kou-Giesbrecht *et al*., 2023), contributing to widespread divergence in future carbon and nitrogen flux simulations across terrestrial biosphere models (Hungate *et al*., 2003; Friedlingstein *et al*., 2014; Zaehle *et al*., 2014; Wieder *et al*., 2015; Meyerholt *et al*., 2020).

Over the past few decades, numerous studies have sought to elucidate plant responses to elevated CO_2_, revealing consistent leaf and whole-plant patterns. At the leaf level, C_3_ plants grown under elevated CO_2_ exhibit increased net photosynthesis rates compared to plants grown under ambient CO_2_ (Medlyn *et al*., 1999; Ainsworth & Long, 2005; Bernacchi *et al*., 2005; Lee *et al*., 2011; Poorter *et al*., 2022). These patterns correspond with reduced mass- and area-based leaf nitrogen content, increased leaf mass per area, reduced stomatal conductance, and reduced photosynthetic capacity, yielding increased photosynthetic nitrogen-use efficiency and water-use efficiency (Curtis, 1996; Drake *et al*., 1997; Medlyn *et al*., 1999; Ainsworth & Long, 2005; Ainsworth & Rogers, 2007; Lee *et al*., 2011; Pastore *et al*., 2019; Poorter *et al*., 2022). At the whole-plant level, C_3_ plants grown under elevated CO_2_ exhibit increased total leaf area, which supports greater net primary productivity and total biomass compared to plants grown under ambient CO_2_ (Coleman *et al*., 1993; Ainsworth *et al*., 2002; Ainsworth & Rogers, 2007; Finzi *et al*., 2007; Poorter *et al*., 2022). Some experiments suggest that elevated CO_2_ increases belowground carbon allocation and the ratio of root biomass to shoot biomass compared to plants grown under ambient CO_2_ (Nie *et al*., 2013), though this allocation response is not consistently observed (Luo *et al*., 1994; Poorter *et al*., 2022).

Despite consistent plant responses to elevated CO_2_ documented across experiments, mechanisms that drive these responses remain unresolved. Some have hypothesized that plant responses to elevated CO_2_ are constrained by nitrogen availability, as net primary productivity is limited by nitrogen availability globally (Vitousek & Howarth, 1991; LeBauer & Treseder, 2008). The progressive nitrogen limitation hypothesis predicts that elevated CO_2_ will increase plant nitrogen uptake to support greater net primary productivity, which will cause nitrogen availability to decline over time (Luo *et al*., 2004). The hypothesis predicts that this response should increase growth and net primary productivity under elevated CO_2_ over short time scales that dampen with time as nitrogen becomes progressively more limiting and stored in longer-lived tissues. Growth responses to elevated CO_2_ expected from the progressive nitrogen limitation hypothesis have received some support from free-air CO_2_ enrichment experiments (Reich *et al*., 2006; Norby *et al*., 2010), though these patterns are not consistently observed (Finzi *et al*., 2006, 2007; Moore *et al*., 2006; Liang *et al*., 2016).

Assuming positive relationships between soil nitrogen availability, leaf nitrogen content, and photosynthetic capacity (Field & Mooney, 1986; Evans, 1989; Evans & Seemann, 1989; Walker *et al*., 2014; Firn *et al*., 2019; Liang *et al*., 2020), the progressive nitrogen limitation hypothesis implies that reductions in nitrogen availability over time might explain why C_3_ plants exhibit decreased leaf nitrogen content and photosynthetic capacity under elevated CO_2_. However, results from free-air CO_2_ enrichment experiments show that reductions in leaf nitrogen content and photosynthetic capacity under elevated CO_2_ are decoupled from changes in nitrogen availability (Crous *et al*., 2010; Lee *et al*., 2011; Pastore *et al*., 2019). Additionally, variance in leaf nitrogen and photosynthetic capacity across environmental gradients tends to be more strongly determined through aboveground growth conditions that set demand to build and maintain photosynthetic enzymes than through changes in soil resource availability (Dong *et al*., 2017, 2020, 2022a; Smith *et al*., 2019; Smith & Keenan, 2020; Paillassa *et al*., 2020; Peng *et al*., 2021; Querejeta *et al*., 2022; Westerband *et al*., 2023; Waring *et al*., 2023). These patterns indicate that leaf photosynthetic responses to elevated CO_2_ may be a product of altered leaf nitrogen demand to build and maintain photosynthetic enzymes and may not be as strongly linked to changes in nitrogen availability.

Eco-evolutionary optimality theory provides a framework for understanding how leaf photosynthetic responses to elevated CO_2_ may be determined through demand to build and maintain photosynthetic enzymes (Harrison *et al*., 2021). Merging photosynthetic least-cost (Wright *et al*., 2003; Prentice *et al*., 2014) and optimal coordination (Chen *et al*., 1993; Maire *et al*., 2012) theories, eco-evolutionary optimality theory posits that reduced leaf nitrogen allocation under elevated CO_2_ is the downstream result of a stronger downregulation in the maximum rate of Ribulose-1,5-bisphosphate (RuBP) carboxylase/oxygenase (Rubisco) carboxylation (*V*_cmax_) than the maximum rate of electron transport for RuBP regeneration (*J*_max_), which reduces leaf nitrogen demand to build and maintain photosynthetic enzymes. Optimal leaf nitrogen allocation to photosynthetic capacity allows plants to make more efficient use of available light while avoiding overinvestment in Rubisco, which has high nitrogen and energetic costs of construction and maintenance (Evans, 1989; Sage, 1994; Evans & Clarke, 2019). Such optimal leaf nitrogen allocation responses to elevated CO_2_ increases photosynthetic nitrogen-use efficiency and allows increased net photosynthesis rates to be achieved through increasingly equal co-limitation of Rubisco carboxylation and electron transport for RuBP regeneration (Chen *et al*., 1993; Maire *et al*., 2012; Wang *et al*., 2017; Smith *et al*., 2019). The expected optimal leaf response to elevated CO_2_ has received some empirical support (Crous *et al*., 2010; Lee *et al*., 2011; Smith & Keenan, 2020; Harrison *et al*., 2021; Dong *et al*., 2022b; Cui *et al*., 2023), though no studies have connected these patterns with concurrently measured whole-plant responses.

The eco-evolutionary optimality hypothesis deviates from the progressive nitrogen limitation hypothesis by indicating that photosynthetic responses to elevated CO_2_ are driven by leaf nitrogen demand to build and maintain photosynthetic enzymes and are independent of changes in soil nitrogen supply. However, the eco-evolutionary optimality hypothesis does not discount the role of soil nitrogen availability on whole-plant responses to elevated CO_2_, where the expected optimal strategy in response to elevated CO_2_ is to allocate surplus nitrogen not needed to satisfy leaf nitrogen demand toward the construction of a greater quantity of optimally coordinated leaves and other plant organs. Thus, whether the supply-driven progressive nitrogen limitation hypothesis or demand-driven eco-evolutionary optimality hypothesis controls plant responses to elevated CO_2_ may be a matter of scale, where leaf photosynthetic responses to elevated CO_2_ are determined through demand to build and maintain photosynthetic enzymes and whole-plant responses to elevated CO_2_ are regulated by changes in nitrogen supply.

Plants allocate carbon belowground in exchange for nutrients through different nutrient acquisition strategies, including direct uptake pathways or symbioses with mycorrhizal fungi and symbiotic nitrogen-fixing bacteria (Gutschick, 1981; Smith & Read, 2008). Carbon costs to acquire nitrogen, or the amount of carbon allocated belowground per unit nitrogen acquired, vary in species with different nitrogen acquisition strategies and are dependent on environmental factors such as atmospheric CO_2_, temperature, light availability, and nutrient availability (Brzostek *et al*., 2014; Terrer *et al*., 2018; Allen *et al*., 2020; Eastman *et al*., 2021; Perkowski *et al*., 2021; Lu *et al*., 2022; Peng *et al*., 2023). Therefore, nitrogen acquisition strategy cannot be ignored when considering effects of nitrogen availability on plant responses to elevated CO_2_. To date, few studies account for acquisition strategy when considering the role of nitrogen availability on leaf and whole-plant responses to elevated CO_2_ (e.g., Terrer *et al*., 2016, 2018; Smith & Keenan, 2020). Such studies found that nitrogen acquisition strategies with reduced carbon costs to acquire nitrogen may buffer the effect of nitrogen limitation at the whole-plant level (Terrer *et al*., 2018), but leaf-level responses remain inconsistent (Terrer *et al*., 2018; Smith & Keenan, 2020).

Here, we conducted a growth chamber experiment using *Glycine max* L. (Merr.) seedlings grown under full factorial combinations of two CO_2_ concentrations, two inoculation treatments, and nine soil nitrogen fertilization treatments to reconcile the role of nitrogen supply and demand on plant responses to elevated CO_2_. We used this experimental setup to test the following hypotheses:

1. Following the demand-driven eco-evolutionary optimality hypothesis, elevated CO_2_ will downregulate *V*_cmax_ more strongly than *J*_max_, increasing *J*_max_:*V*_cmax_ and allowing increased net photosynthesis rates to approach equal co-limitation of Rubisco carboxylation and electron transport for RuBP regeneration. Leaf photosynthetic responses to elevated CO_2_ will be independent of nitrogen fertilization and inoculation treatment and will correspond with increased photosynthetic nitrogen-use efficiency.
2. Following the supply-driven nitrogen limitation hypothesis, positive effects of elevated CO_2_ on total leaf area and total biomass will be enhanced with increasing nitrogen fertilization due to increased plant nitrogen uptake and reduced carbon costs to acquire nitrogen. Inoculation with symbiotic nitrogen-fixing bacteria will enhance positive growth responses to elevated CO_2_, though these responses will only be apparent under low nitrogen fertilization levels where individuals will have increased investment in nitrogen acquisition through symbiotic nitrogen fixation.

## Methods

### Seed treatments and experimental design

*Glycine max* seeds were planted in 144 6-liter surface sterilized pots (NS-600, Nursery Supplies, Orange, CA, USA) containing a steam-sterilized 70:30 volume: volume mix of *Sphagnum* peat moss (Premier Horticulture, Quakertown, PA, USA) to sand (Pavestone, Atlanta, GA, USA). Before planting, all *G. max* seeds were surface sterilized in 2% sodium hypochlorite for 3 minutes, followed by three separate 3-minute washes with ultrapure water (MilliQ 7000; MilliporeSigma, Burlington, MA USA). Subsets of surface-sterilized seeds were inoculated with *Bradyrhizobium japonicum* (Verdesian N-Dure™ Soybean, Cary, NC, USA) in a slurry following manufacturer recommendations (3.12 g inoculant and 241 g ultrapure water per 1 kg seed).

Seventy-two pots were randomly planted with surface-sterilized seeds inoculated with *B. japonicum*, while the remaining 72 pots were planted with surface-sterilized uninoculated seeds. Thirty-six pots in each inoculation treatment were randomly placed in one of two atmospheric CO_2_ treatments (420 and 1000 μmol mol^-1^ CO_2_). Plants in each unique inoculation-by-CO_2_ treatment combination randomly received one of nine nitrogen fertilization treatments equivalent to 0 (0 mM), 35 (2.5 mM), 70 (5 mM), 105 (7.5 mM), 140 (10 mM), 210 (15 mM), 280 (20 mM), 350 (25 mM), or 630 ppm (45 mM) N. Nitrogen fertilization treatments were created using a modified Hoagland’s solution (Hoagland & Arnon, 1950) designed to keep concentrations of all other macronutrients and micronutrients equivalent across treatments (Table S1). Plants received the same nitrogen fertilization treatment twice per week in 150 mL doses as topical agents to the soil surface.

### Growth chamber conditions

Plants were randomly placed in one of six Percival LED-41L2 growth chambers (Percival Scientific Inc., Perry, IA, USA) over two experimental iterations due to chamber space limitation. Two iterations were conducted such that one iteration included all plants grown under elevated CO_2_ plants, and the second iteration included all plants grown under ambient CO_2_. Average (± SD) CO_2_ concentrations across chambers throughout the experiment were 439±5 μmol mol^-1^ CO_2_ for the ambient treatment and 989±4 μmol mol^-1^ CO_2_ for the elevated treatment.

Daytime growth conditions were simulated using a 16-hour photoperiod, with incoming light radiation set to chamber maximum (mean±SD: 1230±12 μmol m^-2^ s^-1^ across chambers), air temperature set to 25°C, and relative humidity set to 50%. The remaining 8-hour period simulated nighttime growing conditions, with incoming light radiation set to 0 μmol m^-2^ s^-1^, chamber temperature set to 17°C, and relative humidity set to 50%. Transitions between daytime and nighttime growing conditions were simulated by ramping incoming light radiation in 45-minute increments and temperature in 90-minute increments over a 3-hour period (Table S2).

Plants grew under average (± SD) daytime light intensity of 1049±27 μmol m^-2^ s^-1^, including ramping periods. In the elevated CO_2_ iteration, plants grew under 24.0±0.2°C during the day, 16.4±0.8°C during the night, and 51.6±0.4% relative humidity. In the ambient CO_2_ iteration, plants grew under 23.9±0.2°C during the day, 16.0±1.4°C during the night, and 50.3±0.2% relative humidity. Within each experiment iteration, any differences in climate conditions across the six chambers were accounted for by shuffling the same group of plants throughout the growth chambers. This process was done by iteratively moving the group of plants on the top rack of a chamber to the bottom rack of the same chamber, while simultaneously moving the group of plants on the bottom rack of a chamber to the top rack of the adjacent chamber. Plants were moved within and across chambers daily during each experiment iteration.

### Leaf gas exchange measurements

Leaf gas exchange measurements were collected on the seventh week of development, before the onset of reproduction. All gas exchange measurements were collected on the center leaf of the most recent fully expanded trifoliate leaf set using LI-6800 portable photosynthesis machines configured with a 6800-01A fluorometer head and 6 cm^2^ aperture (LI-COR Biosciences, Lincoln, NE, USA). Specifically, net photosynthesis (*A*_net_; μmol m^-2^ s^-1^), stomatal conductance (*g*_sw_; mol m^-2^ s^-1^), and intercellular CO_2_ (*C*_i_; μmol mol^-1^) concentrations were measured across a range of atmospheric CO_2_ concentrations (i.e., an *A*_net_/*C*_i_ curve) using the Dynamic Assimilation™ Technique. The Dynamic Assimilation™ Technique corresponds well with traditional steady-state *A*_net_/*C*_i_ curves in *G. max* (Saathoff & Welles, 2021). *A*_net_/*C*_i_ curves were generated along a reference CO_2_ ramp down from 420 µmol mol^-1^ CO_2_ to 20 µmol mol^-1^ CO_2_, followed by a ramp up from 420 µmol mol^-1^ CO_2_ to 1620 µmol mol^-1^ CO_2_ after a 90-second wait period at 420 µmol mol^-1^ CO_2_. The ramp rate for each curve was set to 200 μmol mol^-1^ min^-1^, logging every five seconds, which generated 96 data points per response curve. All *A*_net_/*C*_i_ curves were generated after *A*_net_ and *g*_sw_ stabilized in a LI-6800 cuvette set to a 500 mol s^-1^ flow rate, 10000 rpm mixing fan speed, 1.5 kPa vapor pressure deficit, 25°C leaf temperature, 2000 μmol m^-2^ s^-1^ incoming light radiation, and initial reference CO_2_ set to 420 µmol mol^-1^.

Snapshot *A*_net_ measurements were extracted from each *A*_net_/*C*_i_ curve, both at a common CO_2_ concentration, 420 µmol mol^-1^ CO_2_ (*A*_net,420_; μmol m^-2^ s^-1^), and under each individual’s growth CO_2_ concentration, 420 and 1000 µmol mol^-1^ CO_2_ (*A*_net,growth_; μmol m^-2^ s^-1^). Dark respiration (*R*_d_; μmol m^-2^ s^-1^) measurements were collected with the same leaf used to generate *A*_net_/*C*_i_ curves following at least 30 minutes of darkness. Measurements were collected on a 5-second log interval for 60 seconds after the leaf stabilized in a LI-6800 cuvette set to a 500 mol s^-1^ flow rate, 10000 rpm mixing fan speed, 1.5 kPa vapor pressure deficit, 25°C leaf temperature, and 420 µmol mol^-1^ reference CO_2_ concentration (regardless of CO_2_ treatment), with incoming light radiation set to 0 μmol m^-2^ s^-1^. A single dark respiration value was determined for each leaf by calculating the mean dark respiration value across the logging interval.

### A/C_i_ curve-fitting and parameter estimation

*A*_net_/*C*_i_ curves were fit using the ‘fitaci’ function in the ‘plantecophys’ R package (Duursma, 2015). This function estimates the maximum rate of Rubisco carboxylation (*V*_cmax_; µmol m^-2^ s^-1^) and maximum rate of electron transport for RuBP regeneration (*J*_max_; µmol m^-2^ s^-1^) based on the Farquhar *et al*. (1980) biochemical model of C_3_ photosynthesis. Triose phosphate utilization (TPU) limitation was included as an additional rate-limiting step in all curve fits after visually observing clear TPU limitation for most curves. All curve fits included measured dark respiration values. As *A*_net_/*C*_i_ curves were generated using a common leaf temperature (25°C), curves were fit using Michaelis-Menten coefficients for Rubisco affinity to CO_2_ (*K*_c_; μmol mol^-1^) and O_2_ (*K*_o_; mmol mol^-1^), and the CO_2_ compensation point *(Γ*\*; μmol mol^-1^) reported in Bernacchi *et al*. (2001). Specifically, *K*_c_ was set to 404.9 μmol mol^-1^, *K*_o_ was set to 278.4 μmol mol^-1^, and *Γ*\* was set to 42.75 μmol mol^-1^. For clarity, *V*_cmax_, *J*_max_, and *R*_d_ estimates are referenced throughout the rest of the paper as *V*_cmax25_, *J*_max25_, and *R*_d25_.

### Leaf trait measurements

The leaf used to generate *A*_net_/*C*_i_ curves and dark respiration measurements was harvested immediately following gas exchange measurements. Images of each focal leaf were curated using a flat-bed scanner to determine fresh leaf area using the ’LeafArea’ R package (Katabuchi, 2015), which automates leaf area calculations using ImageJ software (Schneider *et al*., 2012). Post-processed images were visually assessed to check against errors in the automation process. Each leaf was dried at 65°C for at least 48 hours and subsequently weighed and ground until homogenized. Leaf mass per area (*M*_area_; g m^-2^) was calculated as the ratio of dry leaf biomass to fresh leaf area. Leaf nitrogen content (*N*_mass_; gN g^-1^) was quantified using a subsample of ground and homogenized leaf tissue through elemental combustion analysis (Costech-4010, Costech, Inc., Valencia, CA, USA). Leaf nitrogen content per unit leaf area (*N*_area_; gN m^-2^) was calculated by multiplying *N*_mass_ and *M*_area_. Photosynthetic nitrogen-use efficiency (*PNUE*_growth_; µmol CO_2_ g^-1^ N s^-1^) was estimated as the ratio of *A*_net,growth_ to *N*_area_.

Chlorophyll content was extracted from a second leaf in the same trifoliate leaf set as the leaf used to generate *A*_net_/*C*_i_ curves. A cork borer was used to punch between 3-5 0.6 cm^2^ disks from the leaf. Images of each set of leaf disks were curated using a flat-bed scanner to determine wet leaf area, again quantified using the ’LeafArea’ R package (Katabuchi, 2015). Leaf disks were shuttled into a test tube containing 10 mL dimethyl sulfoxide, vortexed, and incubated at 65°C for 120 minutes (Barnes *et al*., 1992). Incubated test tubes were vortexed again before being loaded in 150 μL triplicate aliquots to a 96-well plate. Dimethyl sulfoxide was loaded in each plate as a single 150 μL triplicate aliquot and used as a blank. Absorbance measurements at 649 nm (*A*_649_) and 665 nm (*A*_665_) were recorded in each well using a plate reader (Biotek Synergy H1; Biotek Instruments, Winooski, VT USA), with triplicates averaged and corrected by the mean of the blank absorbance value. Blank-corrected absorbance values were used to estimate *Chl*_a_ (μg mL^-1^) and *Chl*_b_ (μg mL^-1^) following equations from Wellburn (1994):

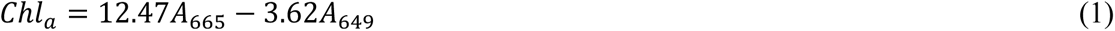

and

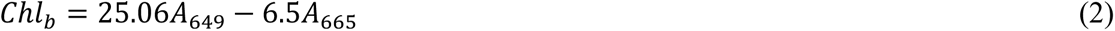

*Chl*_a_ and *Chl*_b_ were converted to mmol mL^-1^ using the molar masses of chlorophyll *a* (893.51 g mol^-1^) and chlorophyll *b* (907.47 g mol^-1^), then added together to calculate the total chlorophyll content in dimethyl sulfoxide extractant (mmol mL^-1^). Total chlorophyll content (mmol) was determined by multiplying the total chlorophyll content in dimethyl sulfoxide by the volume of dimethyl sulfoxide extractant (10 mL). Area-based chlorophyll content (*Chl*_area_; mmol m^-2^) was then calculated by dividing the total chlorophyll content by the total area of the leaf disks.

Subsamples of ground and homogenized leaf tissue were sent to the University of California-Davis Stable Isotope Facility to determine leaf δ^13^C and δ^15^N using an elemental analyzer (Elementar vario MICRO cube elemental analyzer; Elementar Analysensysteme GmbH, Langenselbold, Germany) interfaced to an isotope ratio mass spectrometer (PDZ Europa 20-20 Isotope Ratio Mass Spectrometer, Sercon Ltd., Chestshire, UK). Leaf δ^13^C was used to estimate the time-integrated ratio of leaf intercellular CO_2_ concentration to atmospheric CO_2_ concentration (*χ*, unitless) using leaf δ^13^C and chamber air δ^13^C following Farquhar *et al*. (1989):

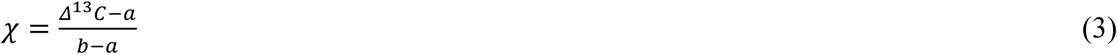

where Δ^13^C represents the relative difference between leaf δ^13^C (‰) and air δ^13^C (‰), and is calculated as:

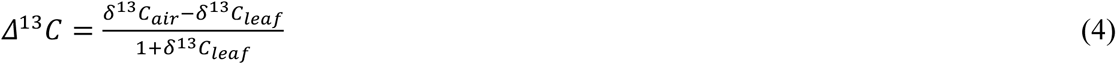

δ^13^C_air_ is the chamber δ^13^C air fractionation, *a* represents the fractionation between ^12^C and ^13^C due to diffusion in air, assumed to be 4.4‰, and *b* represents the fractionation caused by Rubisco carboxylation, assumed to be 27‰ (Farquhar *et al*., 1989). δ^13^C_air_ was quantified in each chamber by collecting air samples in triplicate for each CO_2_ treatment using a 20 mL syringe (Air-Tite Products Co., Inc., Virginia Beach, VA, USA). Each air sample was plunged into a manually evacuated 10 mL Exetainer (Labco Ltd., Lampeter, UK) and sent to the University of California-Davis Stable Isotope Facility, where δ^13^C_air_ was determined using a gas inlet system (GasBenchII; Thermo Fisher Scientific, Waltham, MA, USA) coupled to an isotope ratio mass spectrometer (Thermo Finnigan Delta Plus XL; Thermo Fisher Scientific, Waltham, MA, USA). δ^13^C_air_ for each CO_2_ treatment was estimated by calculating the mean of the triplicate δ^13^C_air_ samples within each chamber, then calculating the mean δ^13^C_air_ across all chambers. Specifically, δ^13^C_air_ was -8.81‰ for the ambient CO_2_ treatment and -5.95‰ for the elevated CO_2_ treatment.

Finally, the percent of leaf nitrogen acquired from the atmosphere (%*N*_dfa_; %) was estimated using leaf δ^15^N and the following equation adapted from Andrews *et al*. (2011):

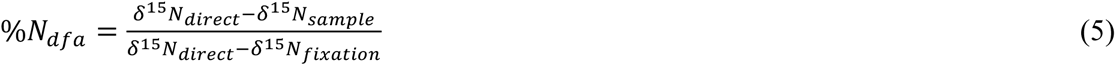

where δ^15^N_direct_ refers to the δ^15^N value from plants that exclusively acquired nitrogen via direct uptake, δ^15^N_sample_ refers to an individual’s leaf δ^15^N, and δ^15^N_fixation_ refers to the δ^15^N value from individuals that were entirely reliant on nitrogen fixation. δ^15^N_direct_ was calculated as the mean leaf δ^15^N of uninoculated individuals within each unique nitrogen fertilization-by-CO_2_ treatment combination. Any individual with visual evidence of root nodule formation or nodule initiation was omitted from the calculation of δ^15^N_direct_. δ^15^N_fixation_ was calculated within each CO_2_ treatment using the mean leaf δ^15^N of inoculated individuals that received 0 ppm N. δ^15^N_fixation_ was not calculated within each unique nitrogen fertilization-by-CO_2_ treatment combination, as previous studies suggest decreased reliance on nitrogen fixation with increasing nitrogen fertilization (e.g., Perkowski *et al*., 2021).

### Whole-plant measurements

Seven weeks after experiment initiation and immediately following gas exchange measurements, all individuals were harvested, and biomass of major organ types (leaves, stems, roots, and nodules when present) were separated. Fresh leaf area of all harvested leaves was measured using a LI-3100C (LI-COR Biosciences, Lincoln, Nebraska, USA). Total fresh leaf area (cm^2^) was calculated as the sum of all leaf areas, including the leaf used to collect gas exchange data and the leaf used to extract chlorophyll content. All harvested material was dried in an oven set to 65°C for at least 48 hours to a constant mass, weighed, and ground to homogeneity. Leaves and root nodules were ground using a mortar and pestle, while stems and roots were ground using an E3300 Single Speed Mini Cutting Mill (Eberbach Corp., MI, USA). Total biomass (g) was calculated as the sum of dry leaf, stem, root, and root nodule biomass. Carbon and nitrogen content was measured for each organ type through elemental combustion (Costech-4010, Costech, Inc., Valencia, CA, USA) using subsamples of ground and homogenized organ tissue. The ratio of root nodule biomass to root biomass was calculated as an additional indicator of investment toward symbiotic nitrogen fixation.

Following Perkowski *et al*. (2021), carbon costs to acquire nitrogen were quantified as the ratio of belowground carbon biomass to total nitrogen biomass (*N*_cost_; gC gN^-1^). Belowground carbon biomass (*C*_bg_; gC) was calculated as the sum of root carbon biomass and root nodule carbon biomass. Root carbon biomass and root nodule carbon biomass were calculated as the product of the organ biomass and respective organ carbon content. Total nitrogen biomass (*N*_wp_; gN) was calculated as the sum of total leaf, stem, root, and root nodule nitrogen biomass. Leaf, stem, root, and root nodule nitrogen biomass was calculated as the product of the organ biomass and respective organ nitrogen content. This calculation does not account for additional carbon costs associated with respiration, root exudation, or root turnover, and therefore may underestimate carbon costs to acquire nitrogen (Perkowski *et al*., 2021).

### Statistical analyses

Uninoculated plants that had substantial root nodule formation (root nodule biomass: root biomass values greater than 0.05 g g^-1^) were removed from analyses under the assumption that plants were either incompletely sterilized or were colonized by symbiotic nitrogen-fixing bacteria from neighboring plants in the chamber. This decision resulted in the removal of sixteen plants from the analysis: two plants in the elevated CO_2_ treatment that received 35 ppm N, three plants in the elevated CO_2_ treatment that received 70 ppm N, one plant in the elevated CO_2_ treatment that received 210 ppm N, two plants in the elevated CO_2_ treatment that received 280 ppm N, two plants in the ambient CO_2_ treatment that received 0 ppm N, three plants in the ambient CO_2_ treatment that received 70 ppm N, two plants in the ambient CO_2_ treatment that received 105 ppm N, and one plant in the ambient CO_2_ treatment that received 280 ppm N.

A series of linear mixed-effects models were built to investigate the impacts of CO_2_ concentration, nitrogen fertilization, and inoculation on *G. max* leaf nitrogen allocation, gas exchange, whole-plant growth, and investment in nitrogen fixation. All models included CO_2_ treatment as a categorical fixed effect, inoculation treatment as a categorical fixed effect, and nitrogen fertilization as a continuous fixed effect, with all possible interaction terms between all three fixed effects also included. Models accounted for climatic differences between chambers across experiment iterations by including a random intercept term that nested the starting chamber rack by CO_2_ treatment. Models with this independent variable structure were created for each of the following dependent variables: *N*_area_, *M*_area_, *N*_mass_, *Chl*_area_, *A*_net,420_, *A*_net,growth_, *V*_cmax25_, *J*_max25_, *J*_max25_:*V*_cmax25_, *R*_d25_, *PNUE_growth_*, *χ*, total leaf area, total biomass, *N*_cost_, *C*_bg_, *N*_wp_, %*N*_dfa_, rood nodule biomass: root biomass, root nodule biomass, and root biomass.

Shapiro-Wilk tests of normality were used to assess whether linear mixed-effects models satisfied residual normality assumptions. All models that did not satisfy residual normality assumptions satisfied such assumptions when response variables were fit using either a natural log or square root data transformation (Shapiro-Wilk: *p*>0.05 in all cases). Specifically, models for *N*_area_, *N*_mass_, *Chl*_area_, *A*_net,420_, *A*_net,growth_, *V*_cmax25_, *J*_max25_, *J*_max25_:*V*_cmax25_, *R*_d25_, *PNUE_growth_*, *χ*, total leaf area, and *N*_cost_ each satisfied residual normality assumptions without data transformation. Models for *M*_area_, total biomass, and *C*_bg_ satisfied residual normality assumptions with a natural log data transformation, while models for *N*_wp_, root nodule biomass: root biomass, root nodule biomass, root biomass, and *%N*_dfa_ satisfied residual normality assumptions with a square root data transformation.

In all models, we used the ‘lmer’ function in the ‘lme4’ R package (Bates *et al*., 2015) to fit each model and the ‘Anova’ function in the ‘car’ R package (Fox & Weisberg, 2019) to calculate Type II Wald’s χ^2^ and determine the significance (*α*=0.05) of each fixed effect coefficient. We used the ‘emmeans’ R package (Lenth, 2019) to conduct post-hoc comparisons using Tukey’s tests, where degrees of freedom were approximated using the Kenward-Roger approach (Kenward & Roger, 1997). Trendlines and error ribbons representing the 95% confidence intervals were drawn in all figures using ‘emmeans’ outputs across the range in nitrogen fertilization values. All analyses and plots were conducted in R version 4.1.0 (R Core Team, 2021). Model results for *χ*, *C*_bg_, *N*_wp_, root nodule biomass: root biomass, root nodule biomass, and root biomass are reported in the *Supplemental Material* (Tables S3-S6; Figs. S3-S6).

## Results

### Leaf nitrogen content

Elevated CO_2_ reduced *N*_area_, *N*_mass_, and *Chl*_area_ by 29%, 50%, and 31%, respectively, and increased *M*_area_ by 44% (*p*<0.001 in all cases; Table 1). Interactions between nitrogen fertilization and CO_2_ (*p*<0.05 in all cases; Table 1) indicated that positive effects of increasing nitrogen fertilization on *N*_area_, *N*_mass_, and *M*_area_ (*p*<0.001 in all cases; Table 1) were stronger under ambient CO_2_ than elevated CO_2_ (Tukey test of the nitrogen fertilization-trait slope between CO_2_: *p*<0.05 in all cases). These responses resulted in a stronger reduction in *N*_area_ and *N*_mass_ and a stronger increase in *M*_area_ under elevated CO_2_ with increasing nitrogen fertilization than ambient CO_2_ (Fig. S1). Nitrogen fertilization did not modify reductions in *Chl*_area_ due to elevated CO_2_ (Tukey test of the nitrogen fertilization-*Chl*_area_ slope between CO_2_ treatments: *p*>0.05).

**Table 1.** Effects of CO_2_ concentration, inoculation, and nitrogen fertilization on leaf nitrogen allocation*.

|  | df | *N*<sub>area</sub> |  | *N*<sub>mass</sub> |  | *M*<sub>area</sub><sup>a</sup> |  | *Chl*<sub>area</sub> |  |
| --- | --- | --- | --- | --- | --- | --- | --- | --- | --- |
| | | $\chi^2$ | *p* | $\chi^2$ | *p* | $\chi^2$ | *p* | $\chi^2$ | *p* |
| CO<sub>2</sub> | 1 | 155.908 | **<0.001** | 272.362 | **<0.001** | 151.319 | **<0.001** | 69.233 | **<0.001** |
| Inoculation (I) | 1 | 86.029 | **<0.001** | 15.576 | **<0.001** | 19.158 | **<0.001** | 136.341 | **<0.001** |
| N fertilization (N) | 1 | 316.408 | **<0.001** | 106.659 | **<0.001** | 21.440 | **<0.001** | 163.111 | **<0.001** |
| CO<sub>2</sub>*I | 1 | 4.729 | **0.030** | 2.025 | 0.155 | 0.029 | 0.866 | 2.102 | 0.147 |
| CO<sub>2</sub>*N | 1 | 5.723 | **0.017** | 22.542 | **<0.001** | 7.619 | **0.006** | 2.999 | 0.083 |
| I*N | 1 | 43.381 | **<0.001** | 11.137 | **0.001** | 5.022 | **0.025** | 75.769 | **<0.001** |
| CO<sub>2</sub>*I*N | 1 | 0.489 | 0.484 | 0.041 | 0.839 | 0.208 | 0.649 | 2.144 | 0.143 |

\*Significance determined using Type II Wald $\chi^2$ tests ( $\alpha=0.05$ ). *P*-values less than 0.05 are in bold. A superscript “a” is included after trait labels to indicate if models were fit with natural log-transformed response variables. Key: df=degrees of freedom, $\chi^2$ =Wald chi- square test statistic, *N*<sub>area</sub>=leaf nitrogen content per unit leaf area (gN m<sup>-2</sup>), *N*<sub>mass</sub>=leaf nitrogen content (gN g<sup>-1</sup>), *M*<sub>area</sub>=leaf mass per unit leaf area (g m<sup>-2</sup>).

An interaction between inoculation and CO_2_ (*p*<0.05; Table 1) indicated that reductions in *N*_area_ due to elevated CO_2_ were stronger in uninoculated plants (36% reduction; Tukey test of the CO_2_ effect in uninoculated plants: *p*<0.001) than inoculated plants (22% reduction; Tukey test of the CO_2_ effect in inoculated plants: *p*<0.001). Inoculation did not modify *N*_mass_, *M*_area_, or *Chl*_area_ responses to elevated CO_2_ (CO_2_-by-inoculation interaction: *p*>0.05 in all cases; Table 1). However, an interaction between nitrogen fertilization and inoculation (*p*<0.05 in all cases; Table 1; Figs. 1a-d) indicated that positive effects of increasing nitrogen fertilization on *N*_area_, *N*_mass_, *M*_area_, and *Chl*_area_ (*p*<0.001 in all cases; Table 1) were stronger in uninoculated plants compared to inoculated plants (Tukey test of the nitrogen fertilization-trait slope between inoculation treatments: *p*<0.05 in all cases).

**Figure 1.**
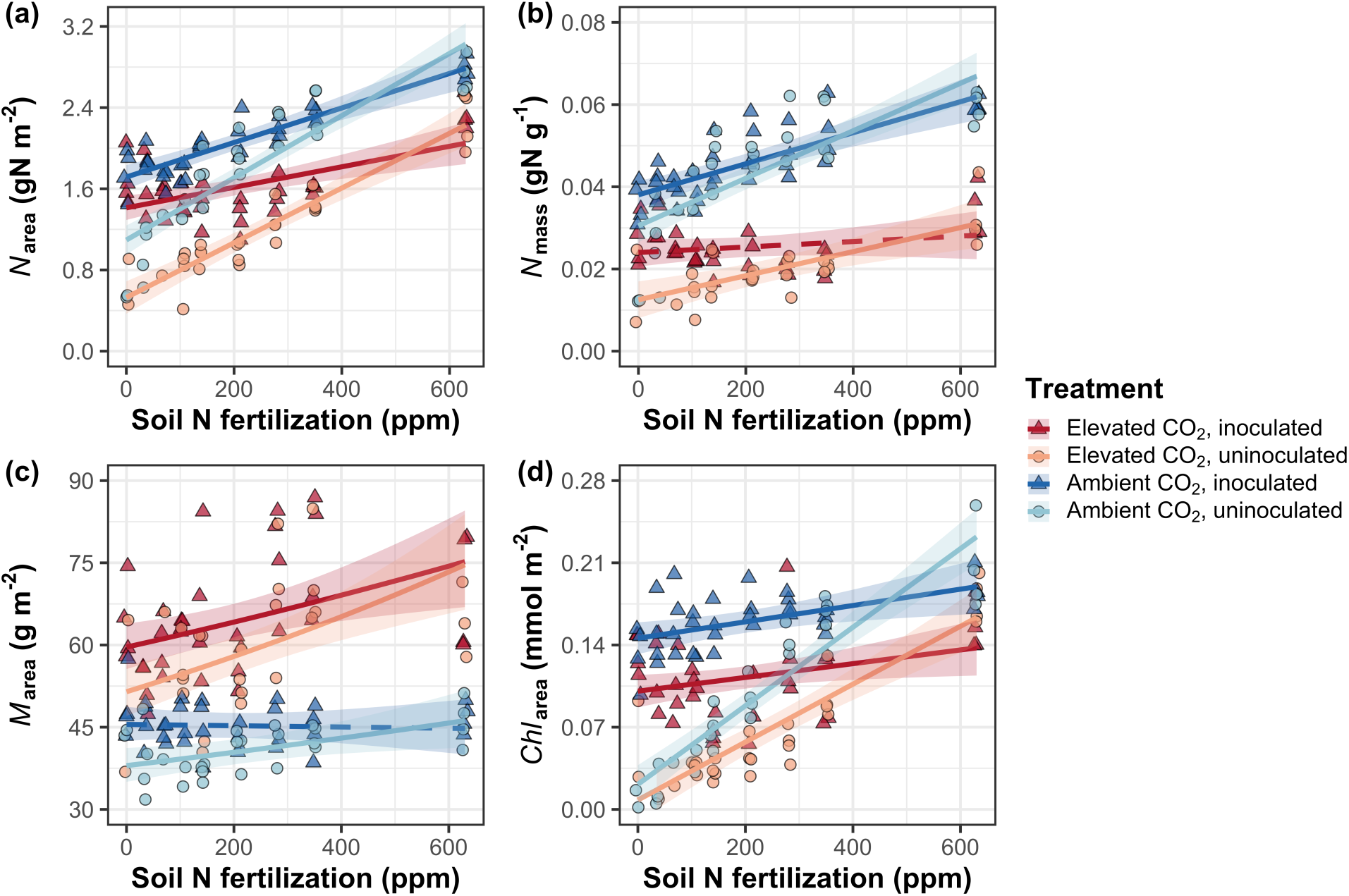
Effects of CO_2_ concentration, nitrogen fertilization, and inoculation on leaf nitrogen per unit leaf area (a), leaf nitrogen per unit leaf mass (b), leaf mass per unit leaf area (c), and chlorophyll content per unit leaf area (d). Nitrogen fertilization is represented on the x-axis in all panels. Red shaded points and trendlines indicate plants grown under elevated CO_2_, while blue shaded points and trendlines indicate plants grown under ambient CO_2_. Light blue and red circular points and trendlines indicate measurements collected from uninoculated plants, while dark blue and red triangular points indicate measurements collected from inoculated plants. Solid trendlines indicate regression slopes that are different from zero (*p*<0.05), while dashed trendlines indicate slopes that are not distinguishable from zero (*p*>0.05).

### Gas exchange

Elevated CO_2_ decreased *A*_net,420_ by 17% (*p*<0.001; Table 2) and increased *A*_net,growth_ by 33% (*p*<0.001; Table 2). Nitrogen fertilization did not modify effects of elevated CO_2_ on *A*_net,420_ or *A*_net,growth_ (CO_2_-by-nitrogen fertilization interaction: *p*>0.05 in both cases; Table 2; Fig. 2a-b). Inoculation did not modify *A*_net,420_ responses to elevated CO_2_ (CO_2_-by-inoculation interaction: *p*>0.05). However, an interaction between CO_2_ and inoculation (*p*<0.05; Table 2) indicated that inoculated plants experienced a stronger increase in *A*_net,growth_ due to elevated CO_2_ (38% increase; Tukey test of the CO_2_ effect in inoculated plants: *p*<0.001) compared to uninoculated plants (26% increase; Tukey test of the CO_2_ effect in uninoculated plants: *p*<0.05). An interaction between nitrogen fertilization and inoculation (*p*<0.001 in both cases; Table 2) indicated that positive effects of increasing nitrogen fertilization on *A*_net,420_ and *A*_net,growth_ (*p*<0.001 in both cases; Table 2; Fig. 2a-b) were stronger in uninoculated plants than inoculated plants (Tukey test comparing the nitrogen fertilization-trait slope between inoculation treatments: *p*<0.001 in both cases).

**Figure 2.**
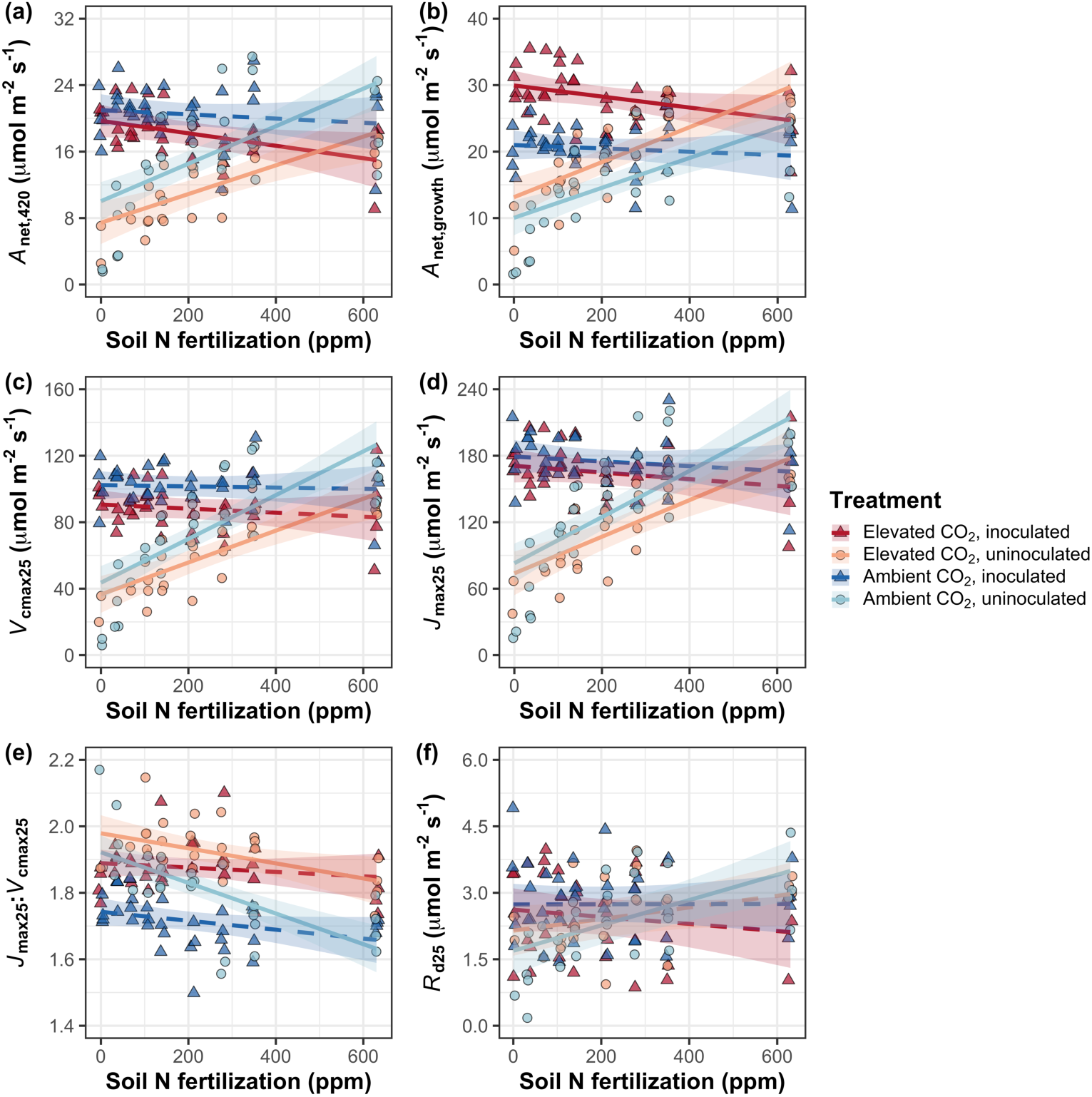
Effects of CO_2_, nitrogen fertilization, and inoculation on net photosynthesis measured at 420 μmol mol^-1^ CO_2_ (a), net photosynthesis measured under growth CO_2_ concentration (b), the maximum rate of Rubisco carboxylation at 25°C (c), the maximum rate of electron transport for RuBP regeneration at 25°C (d), the ratio of the maximum rate of electron transport for RuBP regeneration to the maximum rate of Rubisco carboxylation (e), and dark respiration at 25°C (f). Nitrogen fertilization is represented on the x-axis. Red shaded points and trendlines indicate plants grown under elevated CO_2_, while blue shaded points and trendlines indicate plants grown under ambient CO_2_. Light blue and red circular points and trendlines indicate measurements collected from uninoculated plants, while dark blue and red triangular points indicate measurements collected from inoculated plants. Solid trendlines indicate regression slopes that are different from zero (*p*<0.05), while dashed trendlines indicate slopes that are not distinguishable from zero (*p*>0.05).

**Table 2.**
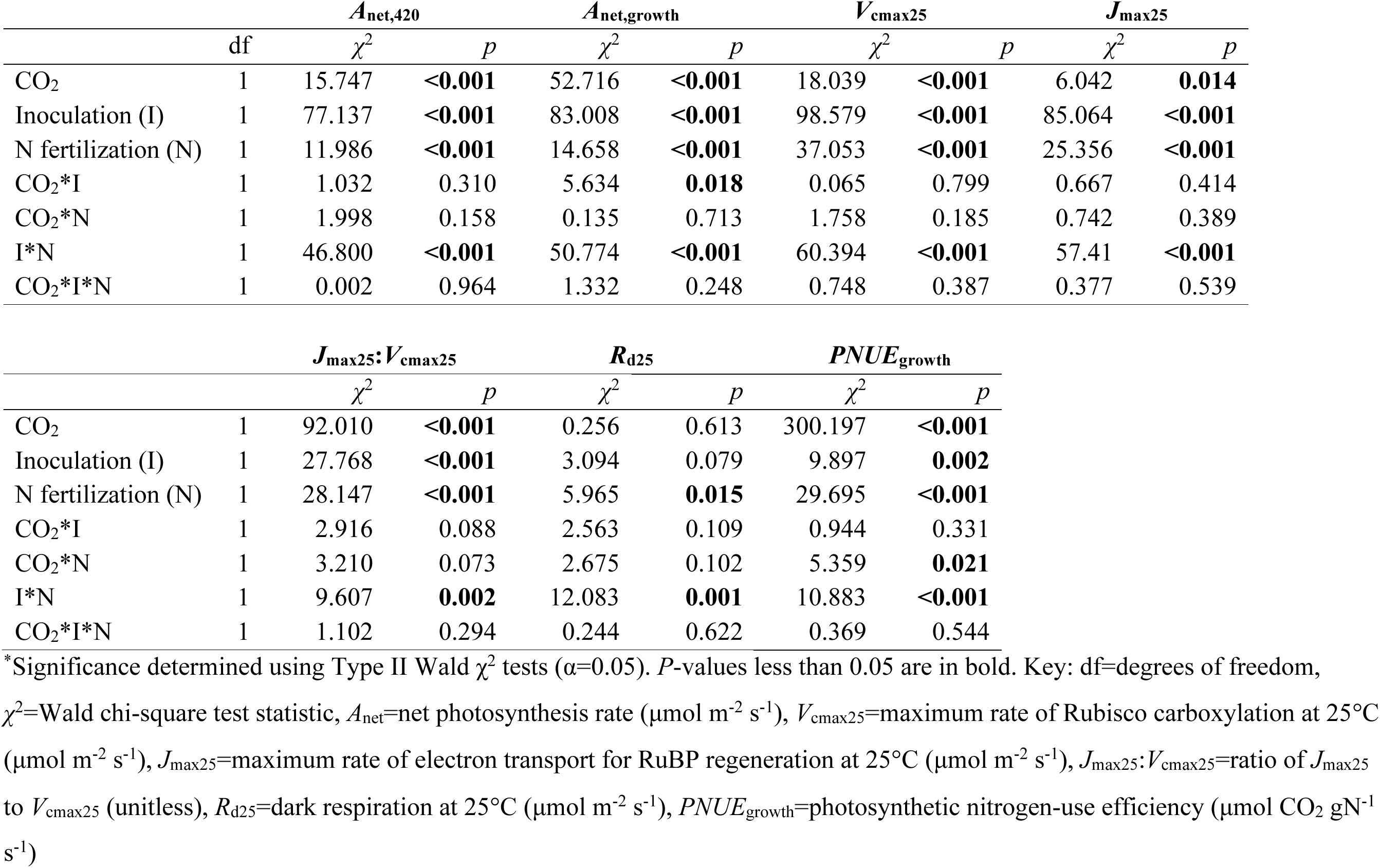
Effects of CO_2_ concentration, inoculation, and nitrogen fertilization on leaf gas exchange*.

Elevated CO_2_ decreased *V*_cmax25_ and *J*_max25_ by 16% and 10%, respectively, increasing *J*_max25_:*V*_cmax25_ by 8% (*p*<0.05 in all cases; Table 2; Fig. 2c-e). *V*_cmax25_, *J*_max25_, and *J*_max25_:*V*_cmax25_ responses to elevated CO_2_ were not modified by nitrogen fertilization (CO_2_-by-nitrogen fertilization interaction: *p*>0.05 in all cases; Table 2; Fig. 2c-e) or inoculation (CO_2_-by-inoculation interaction: *p*>0.05 in all cases; Table 2). An interaction between nitrogen fertilization and inoculation (*p*<0.05 in both cases; Table 2) indicated that positive effects of increasing nitrogen fertilization on *V*_cmax25_ and *J*_max25_ (*p*<0.001 in both cases; Table 2) and negative effects of increasing nitrogen fertilization on *J*_max25_:*V*_cmax25_ (*p*<0.001; Table 2) were driven by uninoculated plants (Tukey test of the nitrogen fertilization-trait slope in uninoculated plants: *p*<0.001 in all cases), as there was no effect of nitrogen fertilization on *V*_cmax25_, *J*_max25_, or *J*_max25_:*V*_cmax25_ in inoculated plants (Tukey test of the nitrogen fertilization-trait slope in inoculated plants: *p*>0.05 in all cases).

There was no effect of CO_2_ concentration on *R*_d25_ (*p*>0.05; Table 2). An interaction between nitrogen fertilization and inoculation (*p*<0.001; Table 2) indicated that the positive effect of increasing nitrogen fertilization on *R*_d25_ (*p*<0.05; Table 2) was driven by uninoculated plants (Tukey test of the nitrogen fertilization-*R*_d25_ slope in uninoculated plants: *p*<0.001), as there was no effect of nitrogen fertilization on *R*_d25_ in inoculated plants (Tukey test of the nitrogen fertilization-*R*_d25_ slope in inoculated plants: *p*>0.05).

### Photosynthetic nitrogen-use efficiency

Elevated CO_2_ increased *PNUE*_growth_ by 90% (*p*<0.001; Table 2; Fig. 3), a pattern that was not modified by inoculation treatment (CO_2_-by-inoculation interaction: *p*>0.05; Table 2). An interaction between CO_2_ and nitrogen fertilization (*p*<0.05; Table 2) indicated that the positive effect of elevated CO_2_ on *PNUE*_growth_ decreased with increasing nitrogen fertilization (Fig. S2). This pattern was driven by a negative effect of increasing nitrogen fertilization on *PNUE*_growth_ (*p*<0.001; Table 2) that was stronger under elevated CO_2_ than ambient CO_2_ (Tukey test comparing the nitrogen fertilization-*PNUE*_growth_ slope between CO_2_ treatments: *p*<0.05). An interaction between nitrogen fertilization and inoculation (*p*<0.001; Table 2; Fig. 3) indicated that the negative effect of increasing nitrogen fertilization on *PNUE*_growth_ was driven by inoculated plants (Tukey test of the nitrogen fertilization-*PNUE*_growth_ slope in inoculated plants: *p*<0.001), as there was no effect of nitrogen fertilization on *PNUE*_growth_ in uninoculated plants (Tukey test of the nitrogen fertilization-*PNUE*_growth_ slope in uninoculated plants: *p*>0.05).

**Figure 3.**
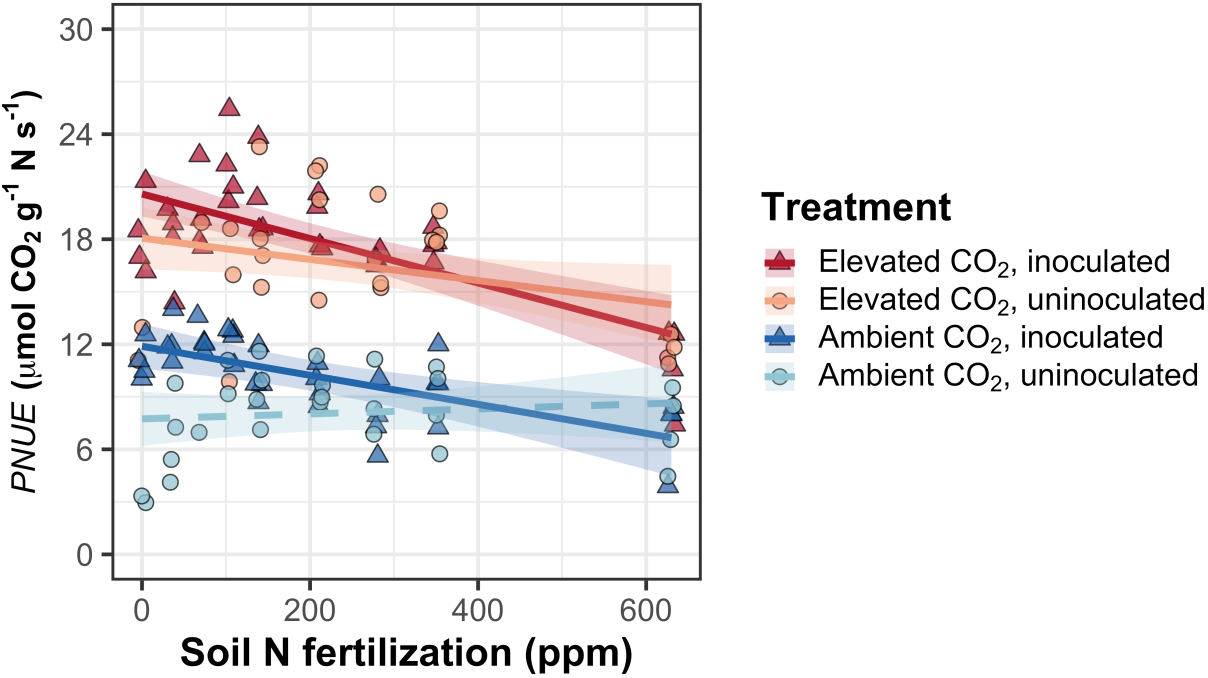
Effects of CO_2_, nitrogen fertilization, and inoculation on photosynthetic nitrogen-use efficiency. Nitrogen fertilization is represented on the x-axis. Red shaded points and trendlines indicate plants grown under elevated CO_2_, while blue shaded points and trendlines indicate plants grown under ambient CO_2_. Light blue and red circular points and trendlines indicate measurements collected from uninoculated plants, while dark blue and red triangular points indicate measurements collected from inoculated plants. Solid trendlines indicate regression slopes that are different from zero (*p*<0.05), while dashed trendlines indicate slopes that are not distinguishable from zero (*p*>0.05).

### Whole-plant traits

Elevated CO_2_ increased total leaf area and total biomass by 51% and 102%, respectively (*p*<0.001 in both cases; Table 3). Positive effects of elevated CO_2_ on total leaf area and total biomass were enhanced with increasing nitrogen fertilization (CO_2_-by-nitrogen fertilization interaction: *p*<0.001 in both cases; Table 3; Fig. 4a-b) but not inoculation (CO_2_-by-inoculation interaction: *p*>0.05 in both cases; Table 3). An interaction between nitrogen fertilization and inoculation (*p*<0.001 in both cases; Table 3) indicated that positive effects of increasing nitrogen fertilization on total leaf area and total biomass (*p*<0.001 in both cases; Table 3) were stronger in uninoculated plants than inoculated plants (Tukey tests comparing the nitrogen fertilization-trait slopes between inoculation treatments: *p*<0.05 for both traits).

**Figure 4.**
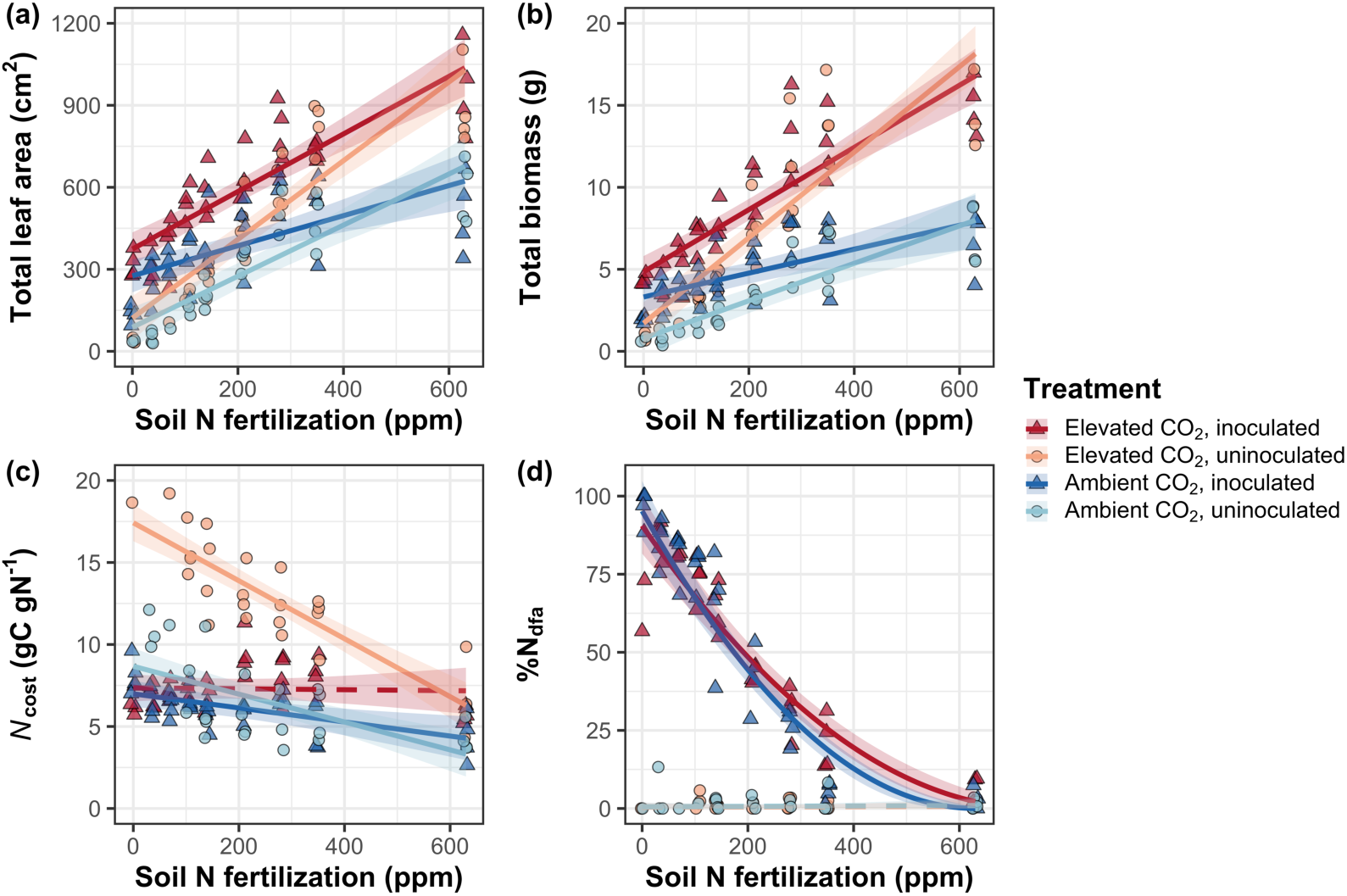
Effects of CO_2_, nitrogen fertilization, and inoculation on total leaf area (a), total biomass (b), structural carbon costs to acquire nitrogen (c), and percent of leaf nitrogen content derived from the atmosphere (d). Nitrogen fertilization is represented on the x-axis. Red shaded points and trendlines indicate plants grown under elevated CO_2_, while blue shaded points and trendlines indicate plants grown under ambient CO_2_. Light blue and red circular points and trendlines indicate measurements collected from uninoculated plants, while dark blue and red triangular points indicate measurements collected from inoculated plants. Solid trendlines indicate regression slopes that are different from zero (*p*<0.05), while dashed trendlines indicate slopes that are not distinguishable from zero (*p*>0.05).

**Table 3.** Effects of CO_2_ concentration, inoculation, and nitrogen fertilization on whole-plant growth, carbon costs to acquire nitrogen, and investment toward symbiotic nitrogen fixation*.

|  | Total leaf area |  |  | Total biomass<sup>b</sup> |  | Carbon cost to<br>acquire nitrogen |  | %N<sub>dfa</sub><sup>b</sup> |  |
| --- | --- | --- | --- | --- | --- | --- | --- | --- | --- |
| | df | $\chi^2$ | *p* | $\chi^2$ | *p* | $\chi^2$ | *p* | $\chi^2$ | *p* |
| CO<sub>2</sub> | 1 | 69.291 | **<0.001** | 131.477 | **<0.001** | 88.189 | **<0.001** | 0.518 | 0.472 |
| Inoculation (I) | 1 | 35.715 | **<0.001** | 34.264 | **<0.001** | 136.343 | **<0.001** | 955.57 | **<0.001** |
| N fertilization (N) | 1 | 274.199 | **<0.001** | 269.046 | **<0.001** | 80.501 | **<0.001** | 292.938 | **<0.001** |
| CO<sub>2</sub>*I | 1 | 2.064 | 0.151 | 0.518 | 0.472 | 85.237 | **<0.001** | 2.010 | 0.156 |
| CO<sub>2</sub>*N | 1 | 18.655 | **<0.001** | 16.877 | **<0.001** | 1.050 | 0.306 | 2.716 | 0.099 |
| I*N | 1 | 10.804 | **0.001** | 15.779 | **<0.001** | 46.489 | **<0.001** | 231.29 | **<0.001** |
| CO<sub>2</sub>*I*N | 1 | <0.001 | 0.990 | 0.023 | 0.880 | 18.125 | **<0.001** | 2.119 | 0.145 |

\*Significance determined using Type II Wald $\chi^2$ tests ( $\alpha=0.05$ ). *P*-values less than 0.05 are in bold and *p*-values between 0.05 and 0.10 are italicized. A superscript “<sup>b</sup>” after trait labels indicates if models were fit using square root transformed variables. Key: df=degrees of freedom, $\chi^2$ =Wald chi-square test statistic, total leaf area (cm<sup>2</sup>), total biomass (g), carbon cost to acquire nitrogen (gC gN<sup>-1</sup>), %N<sub>dfa</sub>=percent leaf nitrogen content fixed from the atmosphere (%).

Elevated CO_2_ increased *N*_cost_ by 62% (*p*<0.001; Table 3), a pattern that was not modified by nitrogen fertilization (CO_2_-by-nitrogen fertilization interaction: *p*>0.05; Table 3). An interaction between CO_2_ and inoculation (*p*<0.05; Table 3) indicated that the positive effect of elevated CO_2_ on *N*_cost_ was stronger in uninoculated plants (99% increase; Tukey test evaluating the CO_2_ effect on *N*_cost_ in uninoculated plants: *p*<0.001) than inoculated plants (21% increase Tukey test evaluating the CO_2_ effect on *N*_cost_ in inoculated plants: *p*<0.05). An interaction between nitrogen fertilization and inoculation (*p*<0.001; Table 3) indicated that the negative effect of increasing nitrogen fertilization on *N*_cost_ (*p*<0.001; Table 3) was stronger in uninoculated plants (Tukey test comparing the nitrogen fertilization-*N*_cost_ slope between inoculation treatments: *p*<0.001). A three-way interaction (*p*<0.001; Table 3) indicated that interactions between nitrogen fertilization and inoculation were stronger under elevated CO_2_ than ambient CO_2_. This pattern was driven by greater *N*_cost_ in uninoculated plants grown under elevated CO_2_ and low nitrogen fertilization than any other CO_2_-by-inoculation treatment combination under low nitrogen fertilization (Tukey test comparing *N*_cost_ in uninoculated individuals grown under elevated CO_2_ and 0 ppm N to all other CO_2_-inoculation treatment combinations grown under 0 ppm N: *p*<0.001 in all cases; Fig. 4c). *N*_cost_ was generally reduced in inoculated plants (*p*<0.001; Table 3). Negative effects of increasing nitrogen fertilization and inoculation on *N*_cost_ were driven by stronger positive effects of each treatment on *N*_wp_ than *C*_bg_, while positive effects of elevated CO_2_ on *N*_cost_ were driven by stronger positive effects on *C*_bg_ than *N*_wp_ (Table S4; Fig. S4).

### Nitrogen fixation

Elevated CO_2_ had no effect on %*N*_dfa_ (*p*=0.472; Table 3; Fig. 4d). An interaction between nitrogen fertilization and inoculation (*p*<0.001; Table 3) indicated that the negative effect of increasing nitrogen fertilization on %*N*_dfa_ (*p*<0.001; Table 3) was driven by inoculated plants (Tukey test of the nitrogen fertilization-%*N*_dfa_ slope in inoculated plants: *p*<0.001), as there was no effect of nitrogen fertilization on %*N*_dfa_ in uninoculated plants (Tukey test of the nitrogen fertilization-%*N*_dfa_ slope in uninoculated plants: *p*>0.05; Fig. 4d).

## Discussion

*Glycine max* seedlings were grown under two CO_2_ concentrations, two inoculation treatments, and nine soil nitrogen fertilization treatments in a full-factorial growth chamber experiment to reconcile the role of nitrogen supply, demand, and acquisition strategy on leaf and whole-plant responses to elevated CO_2_.

Results revealed that elevated CO_2_ increased *A*_net,growth_ despite reduced *N*_area_, *V*_cmax25_, and *J*_max25_. Larger reductions in *V*_cmax25_ than *J*_max25_ increased *J*_max25_:*V*_cmax25_, while respective increases and decreases in *A*_net,growth_ and *N*_area_ increased photosynthetic nitrogen-use efficiency. Effects of elevated CO_2_ on *A*_net,growth_, *V*_cmax25_, *J*_max25_, and *J*_max25_:*V*_cmax25_ were similar across the nitrogen fertilization gradient, suggesting that leaf photosynthetic responses to elevated CO_2_ were decoupled from changes in nitrogen supply. Instead, increased *J*_max25_:*V*_cmax25_ under elevated CO_2_ indicated that plants responded to increasing atmospheric CO_2_ concentrations by allowing enhanced net photosynthesis rates to be achieved by approaching equal co-limitation of Rubisco carboxylation rate-limited photosynthesis and electron transport for RuBP regeneration rate-limited photosynthesis (Chen *et al*., 1993; Maire *et al*., 2012). These responses supported our hypothesis that leaf photosynthetic responses to elevated CO_2_ would be driven by leaf nitrogen demand to build and maintain photosynthetic enzymes and would be independent of nitrogen supply. Leaf photosynthetic responses to elevated CO_2_ corresponded with increased total leaf area and total biomass, patterns that were enhanced with increasing nitrogen fertilization and associated with increased nitrogen uptake efficiency. These results supported our hypothesis that whole-plant responses to elevated CO_2_ would be constrained by nitrogen supply. However, contrasting our hypothesis, inoculation did not modify whole-plant responses to elevated CO_2_ due to similar plant investment in symbiotic nitrogen fixation between CO_2_ treatments.

Combined, results indicate that nitrogen supply and demand were each important factors that determined plant responses to elevated CO_2_ – leaf nitrogen demand to build and maintain photosynthetic enzymes drove leaf photosynthetic responses to elevated CO_2_, while nitrogen supply constrained whole-plant growth responses to elevated CO_2_. These findings support leaf-level patterns expected from eco-evolutionary optimality theory, suggesting that terrestrial biosphere models may improve simulations of leaf photosynthetic processes under future novel environments by considering frameworks that adopt optimality principles (Smith & Keenan, 2020; Harrison *et al*., 2021; Luo *et al*., 2021). Below, we expand and contextualize these conclusions and suggest their implications for terrestrial biosphere model development.

### Nitrogen supply and demand regulate leaf and whole-plant responses to elevated CO_2_ at different scales

Leaf photosynthetic responses to elevated CO_2_ were consistent with previous studies that have investigated or reviewed leaf responses to elevated CO_2_ (Drake *et al*., 1997; Makino *et al*., 1997; Ainsworth *et al*., 2002; Ainsworth & Long, 2005; Ainsworth & Rogers, 2007; Crous *et al*., 2010; Lee *et al*., 2011; Smith & Dukes, 2013; Poorter *et al*., 2022), and follow patterns expected from eco-evolutionary optimality theory (Chen *et al*., 1993; Wright *et al*., 2003; Maire *et al*., 2012; Prentice *et al*., 2014; Wang *et al*., 2017; Smith *et al*., 2019; Smith & Keenan, 2020; Harrison *et al*., 2021). Positive effects of elevated CO_2_ on *A*_net,growth_ and *J*_max25_:*V*_cmax25_ and negative effects of elevated CO_2_ on *V*_cmax25_ and *J*_max25_ were similar across the nitrogen fertilization gradient, indicating that leaf photosynthetic responses to elevated CO_2_ were decoupled from changes in nitrogen supply. Increased *J*_max25_:*V*_cmax25_ and photosynthetic nitrogen-use efficiency under elevated CO_2_ provide strong support for the idea that leaves were downregulating *V*_cmax25_ in response to elevated CO_2_ such that enhanced net photosynthesis rates approached becoming equally co-limited by Rubisco carboxylation and RuBP regeneration (Chen *et al*., 1993; Maire *et al*., 2012; Smith & Keenan, 2020). These patterns suggest that leaf photosynthetic responses to elevated CO_2_ were likely the result of reduced demand to build and maintain photosynthetic enzymes, following patterns expected from eco-evolutionary optimality theory (Harrison *et al*., 2021; Dong *et al*., 2022b).

Whole-plant responses were also consistent with previous studies that have investigated or reviewed whole-plant responses to elevated CO_2_ (Makino *et al*., 1997; Ainsworth *et al*., 2002; Hungate *et al*., 2003; Ainsworth & Long, 2005; Norby *et al*., 2010; Smith & Dukes, 2013; Poorter *et al*., 2022). Greater whole-plant growth under elevated CO_2_ was associated with greater carbon costs to acquire nitrogen through stronger increases in belowground carbon allocation than whole-plant nitrogen uptake. These patterns indicate that plants grown under elevated CO_2_ supported greater total leaf area and total biomass through increased plant nitrogen uptake, though at reduced nitrogen uptake efficiency. Unlike leaf photosynthetic responses to elevated CO_2_, positive whole-plant responses to elevated CO_2_ were enhanced with increasing nitrogen fertilization, supporting our hypothesis that nitrogen supply would constrain whole-plant responses to elevated CO_2_ (Hungate *et al*., 2003; Luo *et al*., 2004; Finzi *et al*., 2007). Positive effects of increasing nitrogen fertilization on total leaf area and total biomass were associated with reductions in carbon costs to acquire nitrogen, a pattern that was driven by stronger increases in whole-plant nitrogen uptake than belowground carbon allocation (Perkowski *et al*., 2021). While reductions in carbon costs to acquire nitrogen due to increasing nitrogen fertilization were similar between CO_2_ treatments, increasing nitrogen fertilization increased whole-plant nitrogen uptake more strongly under elevated CO_2_. This pattern, coupled with similar effects of nitrogen fertilization on belowground carbon allocation responses to elevated CO_2_, indicated that stronger growth responses to elevated CO_2_ with increasing nitrogen fertilization were likely driven by enhanced nitrogen uptake efficiency. These findings suggest that positive short-term effects of nitrogen supply on whole-plant responses to elevated CO_2_ are linked to reduced costs of acquiring nitrogen and increased nitrogen uptake efficiency, supporting conclusions from Terrer *et al*. (2018).

Our findings indicate that nitrogen supply and demand could each explain plant responses to elevated CO_2_, though these factors operated at different scales. Specifically, photosynthetic responses to elevated CO_2_ were determined through reduced leaf nitrogen demand to build and maintain photosynthetic enzymes. Reduced leaf nitrogen demand resulted in a shift in nitrogen allocation to photosynthetic enzymes independent of soil nitrogen supply that increased photosynthetic nitrogen use efficiency and allowed net photosynthesis rates to occur by approaching optimal coordination of Rubisco carboxylation-limited and RuBP regeneration-limited photosynthesis. Whole-plant responses to elevated CO_2_ were enhanced with increasing soil nitrogen supply. Interestingly, optimized nitrogen allocation to photosynthetic capacity may have resulted in nitrogen savings at the leaf level that could have maximized nitrogen allocation to growth. These results suggest that plants grown under elevated CO_2_ responded to increased nitrogen supply by increasing the number of optimally coordinated leaves and that the downregulation in photosynthetic capacity under elevated CO_2_ was not a direct response to changes in nitrogen supply.

### Inoculation with symbiotic nitrogen-fixing bacteria does not modify leaf or whole-plant responses to elevated CO_2_

Inoculation increased *N*_area_, *A*_net,420_, *A*_net,growth_, *V*_cmax25_, *J*_max25_, photosynthetic nitrogen-use efficiency, total leaf area, and total biomass, and decreased *J*_max25_:*V*_cmax25_ and *R*_d25_. These patterns support previous literature suggesting that species that form associations with symbiotic nitrogen-fixing bacteria often have increased leaf nitrogen content, photosynthetic capacity, and growth compared to species that do not form such associations (Adams *et al*., 2016; Bytnerowicz *et al*., 2023). Positive effects of inoculation on leaf and whole-plant traits were strongest under low nitrogen fertilization and rapidly diminished with increasing nitrogen fertilization as investment in symbiotic nitrogen fixation decreased (Andrews *et al*., 2011; Friel & Friesen, 2019; McCulloch & Porder, 2021; Perkowski *et al*., 2021), supporting the idea that nitrogen fixation is a nutrient acquisition strategy that may confer competitive benefits for nitrogen-fixing species growing in low soil nitrogen environments (Rastetter *et al*., 2001; Vitousek *et al*., 2002).

Interestingly, inoculation did not modify effects of elevated CO_2_ on *V*_cmax25_, *J*_max25_, *J*_max25_:*V*_cmax25_, photosynthetic nitrogen-use efficiency, total leaf area, or total biomass. These patterns corresponded with null effects of elevated CO_2_ on %*N*_dfa_ and the ratio of root nodule biomass to root biomass, suggesting that null inoculation effects on plant responses to elevated CO_2_ were primarily due to similar plant investments toward symbiotic nitrogen fixation between CO_2_ treatments. We observed these patterns regardless of nitrogen fertilization level, contrasting our hypothesis that inoculation would enhance whole-plant responses to elevated CO_2_ under low nitrogen fertilization where individuals were expected to be invested more strongly in symbiotic nitrogen fixation. These patterns also contrast previous work showing that inoculated *G. max* is generally more responsive to increasing atmospheric CO_2_ concentrations (Ainsworth *et al*., 2002) and that plant investment toward symbiotic nitrogen fixation tends to be greater under scenarios that increase whole-plant demand to acquire nitrogen (Taylor & Menge, 2018; Friel & Friesen, 2019; McCulloch & Porder, 2021).

### Implications for future model development

Many terrestrial biosphere models predict photosynthetic capacity through parameterized relationships between *N*_area_ and *V*_cmax_ (Rogers, 2014; Rogers *et al*., 2017), which assumes that leaf nitrogen-photosynthesis relationships are constant across growing environments. Our results build on previous work suggesting that leaf nitrogen-photosynthesis relationships dynamically change across growing environments (Smith & Keenan, 2020; Luo *et al*., 2021; Dong *et al*., 2022b; Waring *et al*., 2023), as elevated CO_2_ reduced leaf nitrogen content more strongly than it increased *A*_net,growth_ and decreased *V*_cmax25_ and *J*_max25_. Additionally, positive effects of increasing nitrogen fertilization on indices of photosynthetic capacity were only apparent in uninoculated plants, as there was no effect of nitrogen fertilization on *V*_cmax25_ or *J*_max25_ in inoculated plants. Positive effects of increasing nitrogen fertilization on *N*_area_ and *Chl*_area_ were also markedly weaker in inoculated plants compared to uninoculated plants. These patterns indicate that leaf nitrogen-photosynthesis relationships are context-dependent on nitrogen acquisition strategy, may only be constant in environments where nitrogen supply limits leaf physiology, and will likely shift in response to increasing atmospheric CO_2_ concentrations. Terrestrial biosphere models that predict photosynthetic capacity through parameterized relationships between *N*_area_ and *V*_cmax_ (e.g., Kattge *et al*., 2009; Walker *et al*., 2014) may risk overestimating photosynthetic capacity, therefore net primary productivity and the magnitude of the land carbon sink, under future novel growth environments.

Our results demonstrate that optimal resource allocation to photosynthetic capacity defines leaf photosynthetic responses to elevated CO_2_ and that these responses are independent of nitrogen supply. Current approaches for simulating photosynthetic responses to CO_2_ in terrestrial biosphere models with coupled carbon and nitrogen cycles often invoke patterns expected from progressive nitrogen limitation, where photosynthetic responses to elevated CO_2_ are modeled as a function of positive relationships between nitrogen availability and leaf nitrogen content. Our results contradict this framework, suggesting that photosynthetic responses to elevated CO_2_ are driven by optimal nitrogen investment to satisfy leaf nitrogen demand to build and maintain photosynthetic enzymes. Optimality models that use principles from optimal coordination and photosynthetic least-cost theories (Wang *et al*., 2017; Stocker *et al*., 2020; Scott & Smith, 2022) are capable of capturing responses to CO_2_ independent of nitrogen supply (Smith & Keenan, 2020; Harrison *et al*., 2021), suggesting that including optimality frameworks in terrestrial biosphere models may improve the accuracy by which photosynthetic processes are simulated in response to increasing atmospheric CO_2_ concentrations.

Previous work has highlighted the fact that pot experiments restrict belowground rooting volume and may alter plant allocation responses to environmental change (Ainsworth *et al*., 2002; Poorter *et al*., 2012). In this study, the ratio of pot volume to total biomass was greater under elevated CO_2_ and increased with increasing nitrogen fertilization such that several treatment combinations exceeded values recommended by Poorter *et al*. (2012) to avoid growth limitation imposed by restricted pot volume (<1 g L^-1^; Table S6; Fig. S6). While pot size may have limited plant responses to elevated CO_2_, similar responses to elevated CO_2_ have been observed using field measurements that do not restrict belowground rooting volume (Bernacchi *et al*., 2005; Crous *et al*., 2010; Lee *et al*., 2011; Pastore *et al*., 2019; Smith & Keenan, 2020). Additionally, there was no apparent saturating effect of increasing fertilization on total biomass, belowground carbon biomass, or root biomass under conditions where biomass: pot volume ratios exceeded 1 g L^-1^ (e.g., individuals of either inoculation status grown under high fertilization and elevated CO_2_), which might be expected if pot volume had limited plant growth. The lack of such responses indicate that the pot volume used in this study (6 L) was sufficient to avoid growth limitation.

### Conclusions

Our results indicate that nitrogen supply and demand each helped explain *G. max* responses to elevated CO_2_, though operated at different scales. Supporting eco-evolutionary optimality theory, leaf photosynthetic responses to elevated CO_2_ were independent of soil nitrogen supply and ability to associate with symbiotic nitrogen-fixing bacteria and were instead driven by leaf nitrogen demand to build and maintain photosynthetic enzymes such that net photosynthesis rates approached optimal coordination. Supporting the progressive nitrogen limitation hypothesis, whole-plant responses to elevated CO_2_ were enhanced with increasing nitrogen fertilization due to increased plant nitrogen uptake efficiency coupled with possible cascading effects of nitrogen savings at the leaf level that may have maximized nitrogen allocation to whole-plant growth. However, inoculation did not modify whole-plant responses to elevated CO_2_, as plants invested similarly in symbiotic nitrogen fixation between CO_2_ treatments. Results suggest that plants grown under elevated CO_2_ responded to increased nitrogen supply by increasing the number of optimally coordinated leaves and that the downregulation in photosynthetic capacity under elevated CO_2_ was not modified by changes in nitrogen supply. The differential role of nitrogen supply on leaf and whole-plant responses to elevated CO_2_ coupled with dynamic leaf nitrogen-photosynthesis relationships across CO_2_ and nitrogen fertilization treatments suggests that terrestrial biosphere models may improve simulations of photosynthetic responses to increasing atmospheric CO_2_ concentrations by adopting frameworks that include optimality principles.

## Acknowledgements

This study is a contribution to the LEMONTREE (Land Ecosystem Models based On New Theory, obseRvations and ExperimEnts) project, funded through the generosity of Eric and Wendy Schmidt by recommendation of the Schmidt Futures programme. EAP acknowledges support from a Texas Tech University Doctoral Dissertation Completion Fellowship and a Botanical Society of America Graduate Student Research Award. This work was also supported by US National Science Foundation awards to NGS (DEB-2045968 and DEB-2217353).

## Data Availability

All R scripts, data, and metadata are available at https://doi.org/10.5281/zenodo.10177575 (or on GitHub at: https://github.com/eaperkowski/NxCO2xI_ms_data)

## Author contributions

EAP conceptualized the study objectives and designed the experiment in collaboration with NGS, collected data, conducted data analysis, and wrote the first manuscript draft. EE assisted with data collection and experiment maintenance. NGS conceptualized study objectives and experimental design with EAP and oversaw experiment progress. All authors provided manuscript feedback and approved the manuscript in its current form for submission to *Global Change Biology*.

