## Supplemental Information for "“Nitrogen demand, supply, and acquisition strategy control plant responses to elevated CO_2_ at different scales”"

**Results (cont.)**

*$\chi$*

An interaction between $CO_2$ and nitrogen fertilization ($p<0.001$; Table S3) indicated that the negative effect of increasing nitrogen fertilization on $\chi$ ($p<0.001$; Table S3) was stronger under elevated $CO_2$ than ambient $CO_2$ (Tukey test comparing the nitrogen fertilization-$\chi$ slope between $CO_2$ treatments: $p<0.05$; Fig. S3), resulting in a stronger downregulation of $\chi$ under elevated $CO_2$ with increasing fertilization. A three-way interaction ($p<0.001$; Table S3) indicated that interactions between $CO_2$ and nitrogen fertilization were driven by inoculated plants (Tukey test comparing the nitrogen fertilization-$\chi$ slope between inoculated plants grown under ambient $CO_2$ and inoculated plants grown under elevated $CO_2$: $p<0.001$), as there was no difference in the effect of nitrogen fertilization on $\chi$ between $CO_2$ treatments in uninoculated plants (Tukey test comparing the nitrogen fertilization-$\chi$ slope between uninoculated plants grown under ambient $CO_2$ and uninoculated plants grown under elevated $CO_2$: $p>0.05$). An interaction between $CO_2$ and inoculation ($p<0.001$; Table S3) indicated that elevated $CO_2$ decreased $\chi$ in uninoculated plants (Tukey test of the $CO_2$ effect in uninoculated plants: $p<0.001$) and increased $\chi$ in inoculated plants (Tukey test of the $CO_2$ effect in inoculated plants: $p<0.001$).

*Components of carbon costs to acquire nitrogen*

Elevated $CO_2$ increased $C_{bg}$ by 100% ($p<0.001$; Table S4), a pattern that was not modified by nitrogen fertilization ($CO_2$-by-nitrogen fertilization interaction: $p>0.05$; Table S4). An interaction between $CO_2$ and inoculation ($p<0.05$; Table S4) indicated that the positive effect of inoculation on $C_{bg}$ ($p<0.001$; Table S4) was only apparent under ambient $CO_2$ (Tukey test of the inoculation effect under ambient $CO_2$: $p<0.001$; Fig. S4), as there was no effect of inoculation on $C_{bg}$ under elevated $CO_2$ (Tukey test of the inoculation effect under elevated $CO_2$: $p>0.05$). An interaction between nitrogen fertilization and inoculation ($p<0.001$; Table S3) indicated that the positive effect of increasing nitrogen fertilization on $C_{bg}$ ($p<0.001$; Table S3) was stronger in uninoculated plants than inoculated plants (Tukey test comparing the nitrogen fertilization-$C_{bg}$ slope between inoculation treatments: $p<0.001$).

Elevated $CO_2$ increased $N_{wp}$ by 27% ($p<0.001$; Table S4), a pattern that was enhanced with increasing nitrogen fertilization ($CO_2$-by-nitrogen fertilization interaction: $p<0.05$; Table S4) but was not modified by inoculation ($CO_2$-by-inoculation interaction: $p>0.05$; Table S4). An interaction between nitrogen fertilization and inoculation ($p<0.001$; Table S4) indicated that the positive effect of increasing nitrogen fertilization on $N_{wp}$ ($p<0.001$; Table S4) was stronger in uninoculated plants than inoculated plants (Tukey test comparing the nitrogen fertilization-$N_{wp}$ slope between inoculation treatments: $p<0.001$).

*Nitrogen fixation*

There was no effect of $CO_2$ treatment on root nodule: root biomass ($p>0.05$; Table S5), a pattern that was not modified by nitrogen fertilization ($CO_2$-by-nitrogen fertilization interaction: $p>0.05$; Table S5). However, an interaction between $CO_2$ and inoculation ($p<0.001$; Table S5) indicated that the positive effect of inoculation on root nodule: root biomass ($p<0.001$; Table S5) was stronger under ambient $CO_2$ (3129% increase; Tukey test comparing the inoculation effect under ambient $CO_2$: $p<0.001$) than elevated $CO_2$ (379% increase; Tukey test comparing the inoculation effect under elevated $CO_2$: $p<0.001$). An interaction between nitrogen fertilization and inoculation ($p<0.001$; Table S5) indicated that the negative effect of increasing nitrogen fertilization on root nodule: root biomass ($p<0.001$; Table S5) was stronger in inoculated pots than uninoculated plants (Tukey test comparing the nitrogen fertilization-root nodule: root biomass slope between inoculation treatments: $p<0.001$; Fig. S5).

Root nodule biomass increased by 30% under elevated $CO_2$ ($p<0.001$; Table S5), a pattern that was not modified by nitrogen fertilization ($CO_2$-by-nitrogen fertilization interaction: $p>0.05$; Table S5) or inoculation ($CO_2$-by-inoculation interaction: $p>0.05$; Table S5; Fig. S5). An interaction between nitrogen fertilization and inoculation ($p<0.001$; Table S5) indicated that the negative effect of increasing nitrogen fertilization on root nodule biomass ($p<0.001$; Table S5) was driven by inoculated plants (Tukey test comparing the nitrogen fertilization-root nodule biomass slope in inoculated plants: $p<0.001$), as there was no effect of nitrogen fertilization on root nodule biomass in uninoculated plants (Tukey test comparing the nitrogen fertilization-root nodule biomass slope in uninoculated plants: $p>0.05$; Fig. S5).

Root biomass increased by 96% under elevated $CO_2$ ($p<0.001$; Table S5). An interaction between $CO_2$ concentration and fertilization ($p<0.001$; Table S5) indicated that the positive effect of increasing nitrogen fertilization on root biomass ($p<0.001$; Table S5) was stronger under ambient $CO_2$ (Tukey test comparing the nitrogen fertilization-root biomass slope between $CO_2$ treatments: $p=0.001$). An interaction between $CO_2$ and inoculation ($p<0.001$; Table S5) indicated that the positive effect of inoculation on root biomass ($p<0.001$; Table S5) was driven by the ambient $CO_2$ treatment (Tukey test comparing inoculation effect under ambient $CO_2$: $p<0.001$), as there was no inoculation effect on root biomass under elevated $CO_2$ (Tukey test comparing inoculation effect under elevated $CO_2$: $p>0.05$). An interaction between nitrogen fertilization and inoculation ($p<0.001$; Table S5) indicated that the positive effect of increasing nitrogen fertilization on root biomass ($p<0.001$; Table S5) was stronger in uninoculated plants (Tukey test comparing the fertilization-root biomass slope between inoculation treatments: $p=0.001$).

*The ratio of total biomass to pot volume*

Total biomass: pot volume increased with elevated $CO_2$, inoculation, and nitrogen fertilization ($p<0.001$ in all cases; Table S6). The positive effect of increasing nitrogen fertilization on biomass: pot volume was stronger in uninoculated plants than inoculated plants (Tukey test comparing the nitrogen fertilization-biomass:pot volume slope between inoculation treatments: $p<0.05$; Fig. S6), and when plants were grown under elevated $CO_2$ compared to ambient $CO_2$ (Tukey test comparing the nitrogen fertilization-biomass:pot volume slope between $CO_2$ treatments: $p<0.001$; Fig. S6).

**Table S1** Summary table containing volumes of compounds used to create modified Hoagland's solutions for each soil nitrogen fertilization treatment. All volumes are expressed as milliliters per liter (mL/L)

| **Compound** | **0 ppm N (0 mM N)** | **35 ppm N (2.5 mM N)** | **70 ppm N (5 mM N)** | **105 ppm N (7.5 mM N)** | **140 ppm N (10 mM N)** |
|---|---|---|---|---|---|
| **1 M $NH_4H_2PO_4$** | 0 | 0.165 | 0.33 | 0.5 | 0.67 |
| **2 M $KNO_3$** | 0 | 0.335 | 0.67 | 1 | 1.33 |
| **2 M $Ca(NO_3)_2$** | 0 | 0.335 | 0.67 | 1 | 1.33 |
| **1 M $NH_4NO_3$** | 0 | 0.165 | 0.33 | 0.5 | 0.67 |
| **8 M $NH_4NO_3$** | 0 | 0 | 0 | 0 | 0 |
| **1 M $KH_2PO_4$** | 1 | 0.85 | 0.67 | 0.5 | 0.33 |
| **1 M KCl** | 3 | 2.45 | 2 | 1.5 | 1 |
| **1 M $CaCO_3$** | 4 | 3.33 | 2.67 | 2 | 1.33 |
| **2 M $MgSO_4$** | 1 | 1 | 1 | 1 | 1 |
| **10% Fe-EDTA** | 1 | 1 | 1 | 1 | 1 |
| **Trace elements** | 1 | 1 | 1 | 1 | 1 |

| **Compound** | **210 ppm N (15 mM N)** | **280 ppm N (20 mM N)** | **350 ppm N (25 mM N)** | **630 ppm N (45 mM N)** |
|---|---|---|---|---|
| **1 M $NH_4H_2PO_4$** | 1 | 1 | 1 | 1 |
| **2 M $KNO_3$** | 2 | 2 | 2 | 2 |
| **2 M $Ca(NO_3)_2$** | 2 | 2 | 2 | 2 |
| **1 M $NH_4NO_3$** | 1 | 3.5 | 0 | 0 |
| **8 M $NH_4NO_3$** | 0 | 0 | 0.75 | 2 |
| **1 M $KH_2PO_4$** | 0 | 0 | 0 | 0 |
| **1 M KCl** | 0 | 0 | 0 | 0 |
| **1 M $CaCO_3$** | 0 | 0 | 0 | 0 |
| **2 M $MgSO_4$** | 1 | 1 | 1 | 1 |
| **10% Fe-EDTA** | 1 | 1 | 1 | 1 |
| **Trace elements** | 1 | 1 | 1 | 1 |

**Table S2** Summary of the daily growth chamber growing condition program

| Time | Air temperature (ºC) | PAR ± SD (µmol $m^{-2}$ $s^{-1}$) |
|---|---|---|
| 09:00 | 21 | 278±2 |
| 09:45 | | 557±4 |
| 10:30 | 25 | 797±4 |
| 11:15 | | 1230±12 |
| 22:45 | 21 | 797±4 |
| 23:30 | | 557±4 |
| 00:15 | 17 | 278±2 |
| 01:00 | | 0±0 |

**Table S3** Effects of $CO_2$ concentration, inoculation, and nitrogen fertilization on $\chi$*

| | df | $\chi^2$ | $p$ |
|---|---|---|---|
| $CO_2$ | 1 | 6.809 | **0.009** |
| Inoculation (I) | 1 | 5.827 | **0.016** |
| N fertilization (N) | 1 | 109.544 | **<0.001** |
| $CO_2$*I | 1 | 20.644 | **<0.001** |
| $CO_2$*N | 1 | 11.839 | **<0.001** |
| I*N | 1 | 0.013 | 0.909 |
| $CO_2$*I*N | 1 | 16.901 | **<0.001** |

*Significance determined using Type II Wald $\chi^2$ tests ($\alpha$=0.05). *P*-values less than 0.05 are in bold. Key: df=degrees of freedom, $\chi^2$=Wald chi-square test statistic, $\chi$=isotope-based ratio of intercellular $CO_2$ to extracellular $CO_2$, inversely related to water-use efficiency (unitless)

**Table S4** Effects of $CO_2$ concentration, inoculation, and nitrogen fertilization on components of the carbon cost to acquire nitrogen*

| | | | $\boldsymbol{C_{bg}}$[a] | | $\boldsymbol{N_{wp}}$[b] |
|---|---|---|---|---|---|
| | df | $\chi^2$ | *p* | $\chi^2$ | *p* |
| $CO_2$ | 1 | 84.134 | **<0.001** | 23.890 | **<0.001** |
| Inoculation (I) | 1 | 41.030 | **<0.001** | 134.460 | **<0.001** |
| N fertilization (N) | 1 | 152.248 | **<0.001** | 529.021 | **<0.001** |
| $CO_2$*I | 1 | 8.965 | **0.003** | 1.190 | 0.275 |
| $CO_2$*N | 1 | 1.188 | 0.276 | 5.915 | **0.015** |
| I*N | 1 | 22.648 | **<0.001** | 55.562 | **<0.001** |
| $CO_2$*I*N | 1 | 1.109 | 0.292 | 0.620 | 0.431 |

*Significance determined using Type II Wald $\chi^2$ tests ($\alpha$=0.05). A superscript "[a]" is included after trait labels to indicate if models were fit with natural-log transformed response variables, while a superscript "[b]" is included if models were fit with square-root transformed response variables. *P*-values less than 0.05 are in bold. Key: df=degrees of freedom, $C_{bg}$=belowground carbon biomass (gC, numerator of $N_{cost}$), $N_{wp}$=total nitrogen biomass (gN, denominator of $N_{cost}$).

**Table S5** Effects of $CO_2$ concentration, inoculation, and nitrogen fertilization on investment toward symbiotic nitrogen fixation[*]

| | | **Root nodule biomass[b]** | | **Root biomass[b]** | | **Root nodule: root biomass[b]** | |
|---|---|---|---|---|---|---|---|
| | df | $\chi^2$ | *p* | $\chi^2$ | *p* | $\chi^2$ | *p* |
| $CO_2$ | 1 | 19.258 | **<0.001** | 93.249 | **<0.001** | 0.010 | 0.921 |
| Inoculation (I) | 1 | 755.02 | **<0.001** | 6.983 | **0.008** | 902.063 | **<0.001** |
| N fertilization (N) | 1 | 84.376 | **<0.001** | 195.843 | **<0.001** | 254.741 | **<0.001** |
| $CO_2$*I | 1 | 0.950 | 0.330 | 3.873 | **0.049** | 21.632 | **<0.001** |
| $CO_2$*N | 1 | 2.106 | 0.147 | 11.456 | **<0.001** | 1.590 | 0.207 |
| I*N | 1 | 44.622 | **<0.001** | 7.435 | **0.006** | 132.463 | **<0.001** |
| $CO_2$*I*N | 1 | 0.196 | 0.658 | 0.065 | 0.799 | 2.481 | 0.115 |

[*]Significance determined using Type II Wald $\chi^2$ tests ($\alpha$=0.05). A superscript "[a]" is included after trait labels to indicate if models were fit with natural log-transformed response variables, while a superscript "[b]" is included if models were fit with square-root transformed response variables. *P*-values less than 0.05 are in bold. Key: df=degrees of freedom, root nodule biomass (g), root biomass (g), root nodule: root biomass (unitless).

**Table S6** Effects of $CO_2$ concentration, inoculation, and nitrogen fertilization on the ratio of total biomass to pot volume (g $L^{-1}$)*

| | df | $\chi^2$ | *p* |
|---|---|---|---|
| $CO_2$ | 1 | 146.004 | **<0.001** |
| Inoculation (I) | 1 | 19.320 | **<0.001** |
| N fertilization (N) | 1 | 279.388 | **<0.001** |
| $CO_2$*I | 1 | 0.007 | 0.934 |
| $CO_2$*N | 1 | 49.725 | **<0.001** |
| I*N | 1 | 9.007 | **0.003** |
| $CO_2$*I*N | 1 | 0.640 | 0.434 |

*Significance determined using Type II Wald $\chi^2$ tests (α=0.05). *P*-values less than 0.05 are in bold. Key: df=degrees of freedom, $\chi^2$=Wald chi-square test statistic

### Figure S1

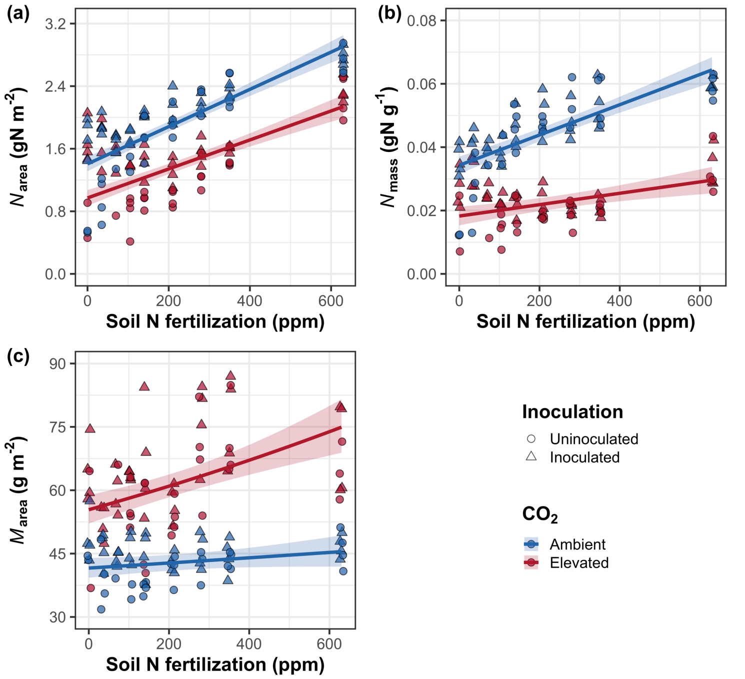


**Figure S1** Effects of $CO_2$ and fertilization inoculation on area-based leaf nitrogen content (a), mass-based leaf nitrogen content (b), and leaf biomass per unit leaf area (c). Fertilization is represented on the x-axis. Red shaded points and trendlines indicate plants grown under elevated $CO_2$, while blue shaded points and trendlines indicate plants grown under ambient $CO_2$. Light blue and red circular points and trendlines indicate measurements collected from uninoculated plants, while dark blue and red triangular points indicate measurements collected from inoculated plants. Solid trendlines indicate regression slopes that are different from zero ($p<0.05$), while dashed trendlines indicate slopes that are not distinguishable from zero ($p>0.05$).

**Figure S2**

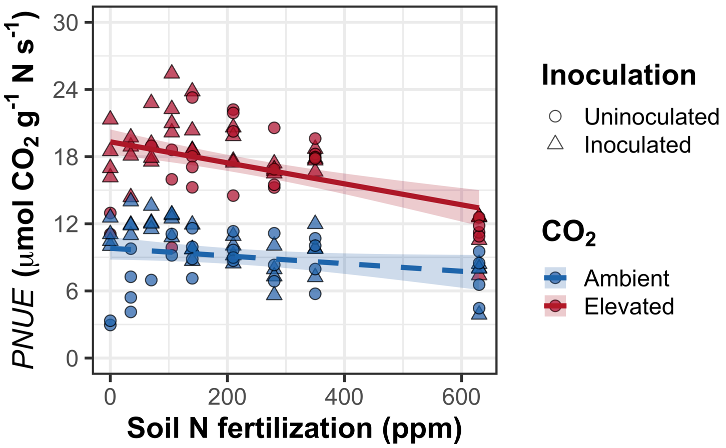


**Figure S2** Effects of $CO_2$ and fertilization inoculation on photosynthetic nitrogen-use efficiency. Fertilization is represented on the x-axis. Red shaded points and trendlines indicate plants grown under elevated $CO_2$, while blue shaded points and trendlines indicate plants grown under ambient $CO_2$. Light blue and red circular points and trendlines indicate measurements collected from uninoculated plants, while dark blue and red triangular points indicate measurements collected from inoculated plants. Solid trendlines indicate regression slopes that are different from zero ($p$<0.05), while dashed trendlines indicate slopes that are not distinguishable from zero ($p$>0.05).

**Figure S3**

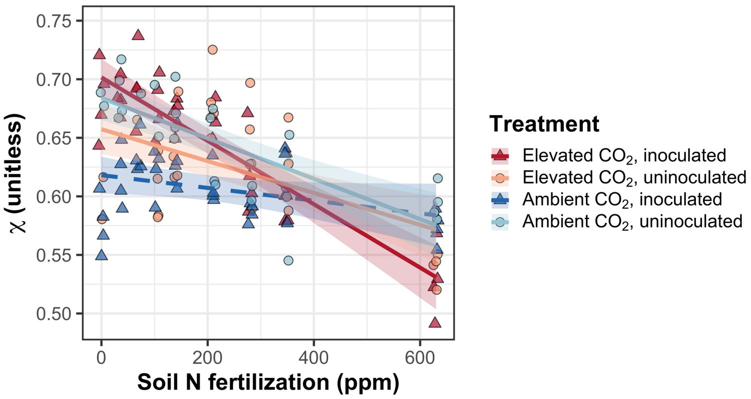


**Figure S3** Effects of nitrogen fertilization, inoculation treatment, and $CO_2$ treatment on $\chi$. Fertilization is represented on the x-axis. Red shaded points and trendlines indicate plants grown under elevated $CO_2$, while blue shaded points and trendlines indicate plants grown under ambient $CO_2$. Light blue and red circular points and trendlines indicate measurements collected from uninoculated plants, while dark blue and red triangular points indicate measurements collected from inoculated plants. Solid trendlines indicate regression slopes that are different from zero ($p$<0.05), while dashed trendlines indicate slopes that are not distinguishable from zero ($p$>0.05).

### Figure S4

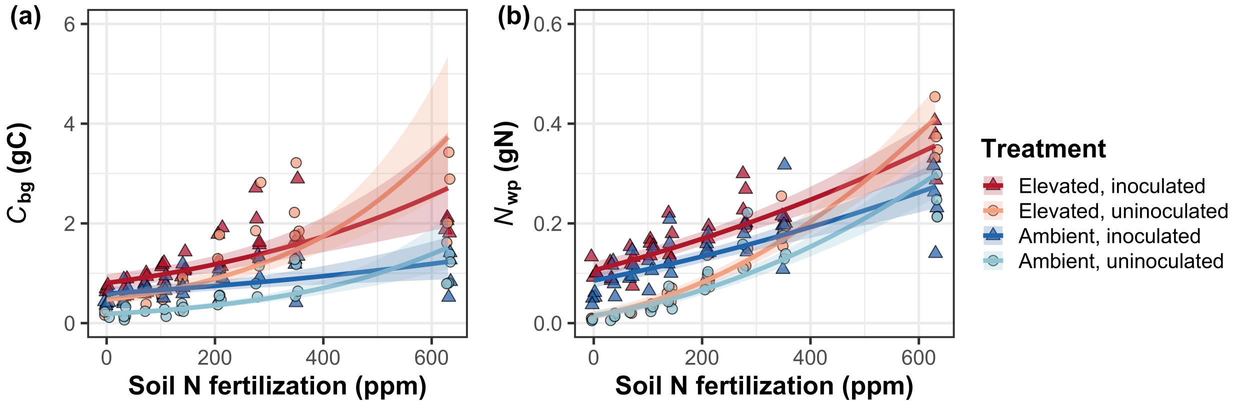


**Figure S4** Effects of $CO_2$ and fertilization inoculation on belowground carbon biomass (a) and total nitrogen biomass (b). Belowground carbon biomass is the numerator of $N_{cost}$, while total nitrogen biomass is the denominator of $N_{cost}$. Fertilization is represented on the x-axis in all panels. Red shaded points and trendlines indicate plants grown under elevated $CO_2$, while blue shaded points and trendlines indicate plants grown under ambient $CO_2$. Light blue and red circular points and trendlines indicate measurements collected from uninoculated plants, while dark blue and red triangular points indicate measurements collected from inoculated plants. Solid trendlines indicate regression slopes that are different from zero ($p$<0.05), while dashed trendlines indicate slopes that are not distinguishable from zero ($p$>0.05).

### Figure S5

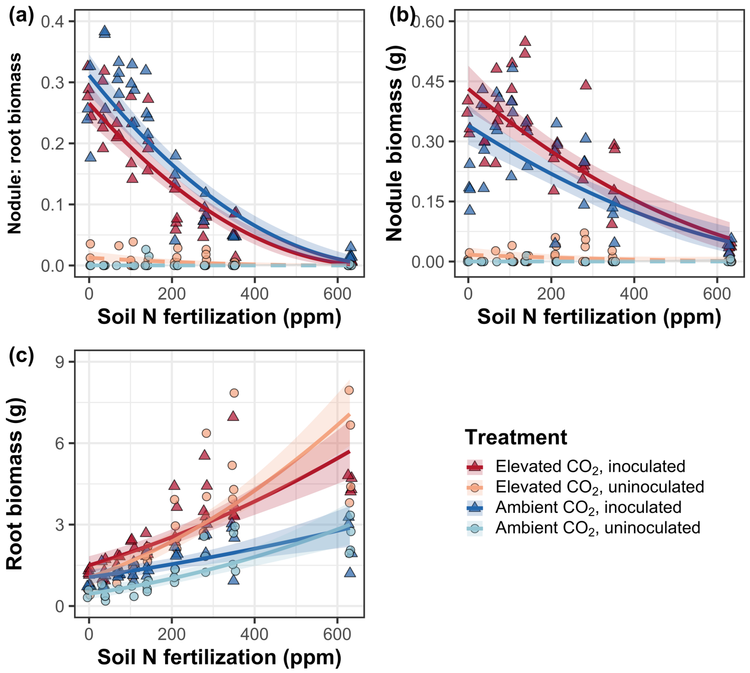


**Figure S5** Effects of nitrogen fertilization, inoculation treatment, and $CO_2$ treatment on root nodule biomass: root biomass (a), root nodule biomass (b), and root biomass (c). Nitrogen fertilization is represented on the x-axis. Red shaded points and trendlines indicate plants grown under elevated $CO_2$, while blue shaded points and trendlines indicate plants grown under ambient $CO_2$. Light blue and red circular points and trendlines indicate measurements collected from uninoculated plants, while dark blue and red triangular points indicate measurements collected from inoculated plants. Solid trendlines indicate regression slopes that are different from zero ($p<0.05$), while dashed trendlines indicate slopes that are not distinguishable from zero ($p>0.05$).

### Figure S6

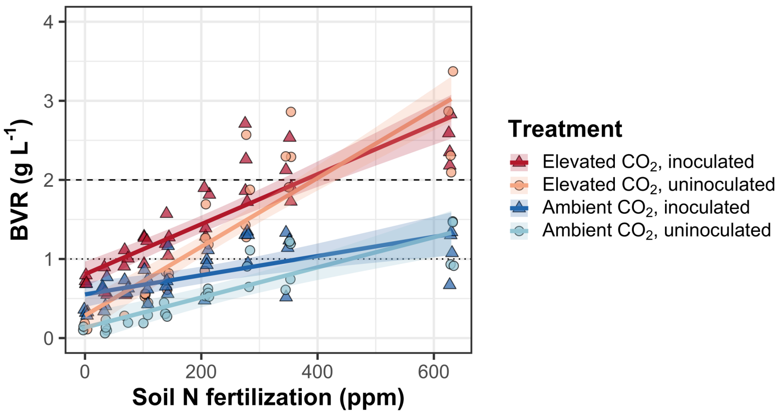


**Figure S6** Effects of $CO_2$, fertilization, and inoculation on the ratio of whole plant biomass to pot volume. Fertilization is represented on the x-axis. Red shaded points and trendlines indicate plants grown under elevated $CO_2$, while blue shaded points and trendlines indicate plants grown under ambient $CO_2$. Light blue and red circular points and trendlines indicate measurements collected from uninoculated plants, while dark blue and red triangular points indicate measurements collected from inoculated plants. Solid trendlines indicate regression slopes that are different from zero ($p$<0.05). The dotted horizontal line indicates the point where biomass: pot volume exceeds 1 g $L^{-1}$, and the dashed line indicates the point where biomass: pot volume exceeds 2 g $L^{-1}$.
